# Competitor Displacement by an Herbivore that Manipulates Plant Defences

**DOI:** 10.1101/2024.05.20.594407

**Authors:** Rachid Chafi, Livia M. S. Ataide, Alessandra Scala, Ernesto Villacis-Perez, Juan M. Alba, Bernardus C. J. Schimmel, Merijn R. Kant

## Abstract

*Tetranychus evansi* is an herbivorous mite specialised on solanaceous hosts, although it has also been observed to colonise non-solanaceous species. It has the ability to suppress the defences of tomato (*Solanum lycopersicum*), and it can displace competitors from this host using a diverse array of traits. *T. evansi* is an invasive species in Africa and Europe, where it often displaces native species. While recent evidence suggests that *T. evansi* can also suppress defences of non-solanaceous hosts, there is a lack of understanding of the molecular changes induced upon mite infestation on hosts other than tomato, as well as how these changes may impact populations of competing herbivores. Here, we investigate the transcriptomic and metabolomic responses of bean (*Phaseolus vulgaris*) to *T. evansi* infestation and to *T. urticae* infestation, a cosmopolitan congeneric that often competes with *T. evansi* for hosts in areas of co-occurrence. We ask whether the presence of *T. evansi* facilitates bean colonization for *T. urticae,* and whether the spatial distribution of mite feeding sites depends on the presence of competitors on bean leaves. We examine whether this facilitation is attributed to jasmonic acid (JA) or salicylic acid (SA) defences by treating plants with exogenous SA and JA and comparing the transcriptomes and metabolomes of bean exposed to either phytohormones or to mite feeding. Finally, we measure phytohormone concentrations and the expression of JA- and SA-responsive genes in plants infested with either mite species or co-infested with both, at different spatial scales. We found that, as previously observed on tomato, *T. urticae* benefits from the suppression of bean defences when sharing a leaf with *T. evansi.* Phytohormone treatments revealed that the reproductive performance of both species decreases with artificially induced JA defences, irrespective of the presence of SA. We found that the molecular suppression and induction of defences is mostly, but not exclusively restricted to the leaf area from where the mites feed. In full leaves co-infested with both mites, levels of marker gene induction were comparable to the inducer mite *T. urticae*, although not as prominent, while in smaller feeding arenas where both species fed closely to each other, the expression of a JA-responsive proteinase inhibitor was suppressed. When residing alone on a leaf, both mites had distinct preferred feeding sites with only partial overlap, but when sharing a leaf, *T. evansi* retained its preferred feeding site and *T. urticae* moved away from its own. We argue that the suppression of defences by *T. evansi* is mostly, although not exclusively, locally restricted, and thus the spatial distribution of individuals on the leaf is a strong determinant of competitor facilitation. This suggests that traits that displace competitors from plant tissues with suppressed defences would be under selection to co-evolve together with defence suppression.

## 1. Introduction

Plants are frequently attacked by herbivores and are therefore under constant pressure to evolve defences that minimize attacks. Some of these defences are constitutively present, such as thick cell walls and cuticles, wax crystals and trichomes, but some defences are inducible upon attack. Inducible defences are produced, released or activated upon the detection of attack signals by the plant, and they are typically regulated by the phytohormones jasmonic acid (JA), salicylic acid (SA), ethylene, among others (Erb & Reymond, 2019). Concomitantly, herbivores are under constant pressure to overcome plant defences and have evolved several different traits to do so. Herbivore adaptations to plant defences include the behavioural avoidance of defended tissues or toxic species (Dussourd, 2017; Kessler et al., 2004; Shroff et al., 2008), the metabolic resistance to toxins due either to target site insensitivity or to the action of detoxification mechanisms (Després et al., 2007; Heckel, 2014; Karageorgi et al., 2019), and the suppression of host defences via the molecular manipulation of plant immunity (Hogenhout & Bos, 2011; Kant et al., 2015; Khan et al., 2018). The evolution of host immunity manipulation is of particular interest for agricultural research, since it can potentially be used for the identification of molecular targets that disrupt the ability of herbivores to tamper with agricultural crops. The suppression of host defences has been in several species of aphids, bugs, gall midges and mites (Blaazer et al., 2018; Bleau et al., 2024; Bunga et al., 2024; Kobayashi, 2016; Zhao et al., 2015). The molecular mechanisms underlying plant defence suppression by herbivores resemble to a certain extent those in plant-pathogen interactions, with the action of effector proteins likely mediating the interaction (Erb & Reymond, 2019; Khan et al., 2018).

Several mite species of the Eriophyidae and Tetranychidae families possess genes encoding for effectors and can suppress plant defences (Greenhalgh et al., 2020; Jonckheere et al., 2018; Jonckheere et al., 2016; R. A. Sarmento et al., 2011; Villarroel et al., 2016). Herbivorous mites are small arthropods that can build large populations in a short time, and thus many species are crop pests (Migeon et al., 2024). Spider mites are cell content feeders that pierce plant cells using their needle-shaped stylets, which inject saliva containing effectors into their host plant prior to ingesting the cell contents (Bensoussan et al., 2016; Blaazer et al., 2018). The suppression of host defences is prevalent in tomato-specialised species such as *Tetranychus evansi*, the tomato red spider mite, and *Aculops lycopersici*, the tomato russet mite (Schimmel et al., 2018). *Tetranychus evansi* is becoming a model system for studies on defence suppression (Alba et al., 2015; Cui et al., 2023; Kant et al., 2008; R. A. Sarmento et al., 2011; B. C. Schimmel et al., 2017; Teodoro-Paulo et al., 2023). Although the suppression trait is always present among *T. evansi* populations, the extent of suppression appears to be variable between native and invasive populations across their geographical range (Knegt et al., 2020). In the extreme generalist species *T. urticae*, the trait is found within populations at low frequencies, while mites that induce defences but that are resistant to them are more prevalent in this species (Alba et al., 2015; Kant et al., 2008). Individuals of both inducing and suppressing species can be susceptible to JA-regulated defences, and thus suppression might be needed for individuals that cannot rely on detoxification mechanisms (Ataide et al., 2016; Renato A Sarmento et al., 2011; B. C. Schimmel et al., 2017; Schimmel et al., 2018; Zhurov et al., 2014). In experimental evolution assays, suppression evolves as an adaptive trait in tomato-selected populations of *T. urticae* (Bruinsma et al., 2023; Wybouw et al., 2015). Together, these observations suggest that suppression is under selection in the field, and that it can evolve in mite populations as an adaptation to overcome host defences. However, in ecologically complex communities, suppression can be prone to exploitation, since the impaired defences of a plant can open up a previously unavailable niche to competitors that cannot suppress defences (Blaazer et al., 2018; Fragata et al., 2022; Glas et al., 2014; R. A. Sarmento et al., 2011).

Interestingly, the suppression of tomato defences by either *T. evansi* or by *A. lycopercisi* benefits *T. urticae* in experiments under controlled conditions, but in nature, *T. evansi* displaces competing native mite species such as *T. urticae* from solanaceous hosts (Boubou et al., 2011; Ferragut et al., 2013; Renato A Sarmento et al., 2011; Schimmel et al., 2018). This incongruency may result from several different factors. *Tetranychus evansi* likely originated in South America, and then spread as an invasive species to Africa and the Mediterranean basin in Europe; areas with climate conditions that can benefit defence-suppressing species (Boubou et al., 2011; Teodoro-Paulo et al., 2024). Outside of its native range, the lack of adapted predators and entomopathogenic fungi can help *T. evansi* build large populations and thus outcompete native mite species (Ferragut et al., 2013). *T. evansi* produces a very dense web that prevents competitors or predators to accessing its feeding sites (Renato A Sarmento et al., 2011). In addition, reproductive interference with genetically incompatible congenerics and the overcompensation of oviposition in the presence of competitors aid *T. evansi* to increase its competitive potential (Sato & Alba, 2020; Sato et al., 2014). Whether these ‘buffering traits’ have evolved along the ability to suppress plant defences in order to protect it from exploitation remains an unexplored possibility that might explain the observed patterns of competitor displacement by suppressing mites (Blaazer et al., 2018).

Whether competitors can exploit a plant with impaired defences will largely depend on how local vs. systemic is the effect of suppression across plant tissues, as well as on the identity of the exploiter. *T. urticae* induces the molecular upregulation of defences across different host species (Agut et al., 2015; Díaz-Riquelme et al., 2016; He et al., 2020; Leitner et al., 2005; Maserti et al., 2011; Zhang et al., 2019; Zhurov et al., 2014). Defence induction by *T. urticae* feeding prompts a change in the tomato transcriptome, in which biological processes such as primary metabolism, growth and photosynthesis processes are downregulated, while processes related to jasmonic acid (JA) signalling, defence responses, amino acid production and specialised metabolite production are induced (Alba et al., 2015; Bruinsma et al., 2023; Wybouw et al., 2015). In contrast, feeding by the tomato specialist *T. evansi* is characterised by JA-signalling and defence responses that are much more attenuated compared to *T. urticae*, as well as by the suppression of biological processes related to the production of specialised metabolites and an even stronger suppression of plant growth (Bruinsma et al., 2023). While this transcriptomic reconfiguration seems to be mostly locally restricted to the area of mite feeding, evidence suggests that these effects can affect adjacent tissue of the same leaf, as well as systemic effects on leaves adjacent to those attacked, and that these systemic effects impact competitor fitness (Bruinsma et al., 2023; R. A. Sarmento et al., 2011; B. C. Schimmel et al., 2017). For example, leaf areas attacked by defence-inducing *T. urticae* and systemic leaves within the attacked plant lower the fitness of competing *T. evansi*, which is susceptible to tomato defences that are already mounted on the attacked tissue (R. A. Sarmento et al., 2011). Leaves attacked by defence-suppressing *T. evansi* benefit both mite species when sharing the same leaf, and *T. evansi* can also benefit from the defence suppression of congenerics in undamaged, systemic leaves (Bruinsma et al., 2023). In addition, *T. evansi* can enhance its reproductive performance when sharing a leaf with competitors, allegedly to increase its competing capacity (B. C. Schimmel et al., 2017). These observations suggest that mites perceive the different physiological changes of an attacked tomato plant, but whether mites modify their behaviour (e.g., the choice of feeding site or feeding intensity) to respond to these molecular changes is poorly understood, particularly at small spatial scales within plants, where these interactions occur. Despite *T. evansi* being able to colonise over 30 plant species outside of the Solanaceae (Migeon et al., 2024), and it being able to suppress defences of a number of these hosts (Paulo et al., 2018), there is a lack of knowledge of the molecular and metabolic changes that *T. evansi* elicits on hosts beyond tomato, as well as whether competitor interactions and behaviour are affected by the molecular changes caused by mite feeding.

In this study, we aim to characterise the molecular mechanisms and behavioural consequences of defence suppression on bean (*Phaseolus vulgaris*), a widespread host within the feeding range of both *T. urticae* and *T. evansi* (Boom et al., 2003; Tahmasebi et al., 2014). Fabaceae and Solanaceae make use of markedly different defences (Ahuja et al., 2012; Wink, 2003), but evidence suggests that the production of proteinase inhibitors, which can affect mite nutrient acquisition, is impaired on bean plants fed on by *T. evansi* (Bruinsma et al., 2023). To achieve our aim, we first establish whether defence-inducing *T. urticae* benefits from the presence of defence-suppressing *T. evansi* on co-infested bean leaves. We then quantify the transcriptomic and metabolic responses of bean plants infested with either *T. evansi* or *T. urticae* and compare them to treatments with exogenous phytohormone application. We also quantify mite fitness on plants and arenas treated with phytohormones to establish the susceptibility of each mite species to bean defences. We further quantify the expression of marker genes and the accumulation of phytohormones in treatments either with single-species infestations or on leaves co-infested with both mite species. To understand the spatial extent of defence suppression, we quantify marker gene expression at two different spatial scales within bean leaves co-infested with both mite species: when they are either physically separated to different areas of the leaf, and when they are restricted to a small leaf area and are thus forced to feed closely to each other. Finally, we investigate the spatial distribution of individuals of both species in single-species infestations on bean leaves and in co-infestations with both species, to understand whether the presence of competitors on the leaf alters the choice of feeding site of each species.

## 2. Material and Methods

### 2.1. Plants and mites

Common bean (*Phaseolus vulgaris* L. cv. Speedy) seeds were sown in potting soil (“potgrond nr. 3”; Jongkind B.V.) and grown in a greenhouse at 25°C: 18°C, 16h L: 8h (day:night photoperiod) and 50-60% relative humidity (RH)) for 10 days before they were transferred to a climate room under standard conditions (25°C, 16L:8D photoperiod, 60% RH). All experiments involving plants were carried out with intact 12-day-old bean plants in a climate room and one of the two first true leaves was used for treatments or mite infestations.

### 2.2. Spider mites

*Tetranychus urticae* Koch ‘Santpoort-2’ (Alba et al., 2015), hereon referred to as “Tu”, was reared on detached bean leaves. *T. evansi* Baker & Pritchard ‘Viçosa-1’ (Alba et al., 2015), hereon referred to as “Te”, was transferred from a previously established rearing on *Solanum lycopersicum* cv. Castlemart to a rearing on detached bean leaves, at least one month before the start of experiments. Spider mite rearings were kept in a climate room under standard conditions. For all plant infestation experiments and mite performance assays, 2 to 4-day-old adult female mites were obtained by creating age-synchronized cohorts two weeks before the experiments.

### 2.3. Reproductive performance of T. urticae on leaves shared with T. evansi

To assess whether the performance of *T. urticae* is affected by the presence of *T. evansi* on a leaf, we carried out a co-infestation assay following Alba *et al*. (2015). Briefly, 1:1 (v/v) mix of insect glue (Cola Entomologica Bio-Controle, São Paulo, Brazil) and lanolin (Sigma-Aldrich, St Louis, MO, USA) was applied around the petiole and middle of the leaf (perpendicular to the midrib) as a barrier to isolate two leaf areas. We created three treatments to assess the number of eggs laid by adult *T. urticae* females, each consisting of five adult *T. urticae* females placed at tip of the leaf and either 15 adult *T. evansi* (15 Te + 5 Tu) or 15 adult *T. urticae* females (15 Tu + 5 Tu) placed on the base of the leaf, or with the base without mites (un-infested + 5 Tu). Two days later, the number of alive mites as well as the number of eggs laid at the tip section were counted using a stereomicroscope. Per treatment, 20-23 plants were infested with mites. Leaves from which all *T. urticae* had migrated were excluded from the analysis. Reproductive performance per female per day was assessed using a linear model with treatment as fixed factor using the command *aov* in R (v 3.5.1).

### 2.4. Reproductive performance of mites feeding on phytohormone-treated leaves

To assess whether mites were susceptible or resistant to bean defences, we assessed the reproductive performance of *T. urticae* and *T. evansi* females feeding on plants treated with the phytohormones JA and SA. To do so, we conducted two independent experiments to account for a possible effect of two commonly used experimental set ups, one using intact plants and one using small discs cut out from bean leaves (Kant et al. 2004, Sarmento et al. 2011, Alba et al. 2015). Using a needleless syringe, we infiltrated the abaxial side of one leaf per bean plant with either 100 µM JA + 100 µM Ile (JA treatment), with 100 µM SA (SA treatment), or with a control solution containing an equivalent amount of methanol (Mock treatment). One day later, five adult female mites of each species were transferred to each infiltrated leaf. After four days of infestation, the numbers of alive mites and eggs laid were counted, samples without surviving mites were excluded. We conducted the assay in two experimental blocks, with a total of n = 16 plants per treatment. In a separate assay, we infiltrated bean leaves of intact plants using the procedure described above, but with either 50 µM JA with 50 µM Ile (JA treatment), 50 µM SA (SA treatment), a combination of the JA and SA treatments (JA + SA treatment), or a mock solution (Mock treatment). One day later, leaf discs (Ø 15 mm) were punched from the infiltrated area, placed on a bed of wet cotton, and a single adult female mite was transferred to each leaf disc. After three days, the numbers of alive mites and eggs laid were counted. Only leaf discs with alive mites were used for analysis. The assay was conducted in two experimental blocks with a total of n = 45 leaf discs per treatment. We also compared the proportion of missing mites, i.e. the mites that died, got stuck, or went missing during the experiment. Differences in reproductive performance per female per day and in the percentage of missing mites were analysed using linear mixed-effects models, with treatment as fixed factor and experimental block as a random factor, using the command *lmer* in package lme4 (R v 3.5.1). Post hoc analyses between treatments were performed using a Tukey corrected p-value with the command *TukeyHSD*.

### 2.5 Transcriptomic analysis of hormone-treated and mite-infested bean leaves

To characterise the transcriptomic changes of bean in response to mite feeding, we sequenced the entire transcriptome of leaves exogenously treated with JA, with SA or infested with mites. Specifically, we created five treatments: 1) intact bean leaves infiltrated with JA [250 µM] or with 2) SA [250 µM], plants infested with either 3) 30 *T. urticae* or with 4) 30 *T. evansi* adult females and 5) clean plants as control. Treated leaves were collected 12 hours post infiltration for the phytohormone treatments, and after 4 days post mite infestation along with their respective controls. Per replicate, the treatments were timed such that all samples were harvested at the same moment in time during the light period. The plants for the five treatments were sown, grown and harvested at the same time. Three biological replicates were collected per treatment (n = 3). Total RNA was extracted using E.Z.N.A.(R) Plant RNA Kit (Omega bio-tek) according to the manufacturer’s instructions. Total RNA was combined with ERCC RNA Spike-In Mix 1 (Thermo Fisher Scientific) and a poly-A enrichment was performed using the Dynabeads mRNA DIRECT Purification Kit (Thermo Fisher Scientific). Bar-coded RNA libraries were generated according to the manufacturer’s protocols using the Ion Total RNA-Seq Kit v2 and the Ion Xpress RNA-Seq barcoding kit (Thermo Fisher Scientific). The size distribution and yield of barcoded libraries were assessed using a 2200 TapeStation System with Agilent D1000 ScreenTapes (Agilent Technologies). Sequencing templates were prepared on an Ion Chef System using an Ion PI Hi-Q Chef Kit (Thermo Fisher Scientific). Strand specific sequencing was performed on an Ion Proton System using an Ion PI v3 chip (Thermo Fisher Scientific) according to the instructions of the manufacturer. After quality control, short reads (<25nt) were discarded. The reads were subsequently mapped on the ERCC spikes using Bowtie2 (Langmead and Salzberg, 2012). Unmapped reads were then mapped on the *P. vulgaris* transcriptome (Phaseolus vulgaris v2.1) with Tmap (http://github.com/iontorrent/tmap). The bean loci were originally annotated using *Arabidopsis thaliana* as a reference (Schmutz et al., 2014). We additionally annotated all genes using the genome of tomato *Solanum lycopersicum* (ITAG 3.2)(Sato et al., 2012) as a reference via blastx with a bitscore of 1e-10 or lower. For our analyses we only included those loci for which at least 10 reads had been mapped in one of the 15 samples. We also removed the reverse compliment RNA from the dataset. Loci for which reads had been mapped in only one of the two treatments were marked “UP” or “DOWN”. Loci for which no reads had been mapped for either treatment were marked as “NA”. Genes that were not annotated were also included in the analysis of differentially regulated genes. Transcript amounts were log(x+0,1) transformed prior to performing statistics. To evaluate differences in gene expression between pairs of treatments we performed a paired t-test and used corrected p-values using a false discovery rate (FDR) of 5% (Benjamini & Hochberg, 1995).

### 2.6. Untargeted metabolomic analysis of hormone-treated and mite-infested bean leaves

In an independent experiment, we sampled bean leaves to conduct an untargeted metabolomics screening of phytohormone-treated and mite-infested leaves using a LC-MS. Samples were obtained following the experimental set up described for RNAseq (see Section 2.5). These were bean leaves infiltrated with either 250 µM JA or 250 µM SA (collected 12 hours post infiltration), and leaves infested with either 30 *T. urticae* or 30 *T. evansi* adult females (collected 4 days post infestation). Leaves from untreated, un-infested plants were sampled as controls. Four biological replicates were collected per treatment (n = 4) and kept at -80C. Mass spectrometry-based metabolomics and subsequent data analysis was performed by MetaSysX GmbH (Potsdam, Germany). Samples were prepared according to MetaSysX standard procedures following a modified protocol from (Salem et al., 2016). Approximately 15 mg of frozen plant material was ground into a fine powder using a tissue homogenizer (Precellys 24, Bertin Technologies, Aixen-Provence, France). The polar metabolites and lipophilic compounds were extracted by a methyl-tert-butyl-ether (MTBE)/methanol/water solvent system, which separates molecules into aqueous and organic phase, respectively. After MTBE extraction, the organic phase containing lipids and lipophilic compounds was transferred to a new 1.5 ml tube. The leftover of the organic phase was removed with a vacuum aspirator and then 500 µl of the aqueous solution containing semi-polar and polar compounds were transferred to a new tube. All the samples were dried using a centrifugal evaporator. The dried samples were resuspended in 170 µl of water or acetonitrile for polar and lipid measurements, respectively.

#### 2.6.2. LC-MS measurements

Metabolites were separated by ultraperformance liquid chromatography (UPLC) (Waters, ACQUITY) and analyzed on a Q Exactive mass spectrometer (Thermo Fisher Scientific) in positive and negative ionization modes. A 2 µl injection volume was analyzed. C8 (100mm × 2.1mm × 1.7μm particles; Waters) and HSS T3 C18 (100mm × 2.1mm × 1.8μm particles; Waters) columns were used for the lipophilic and the hydrophilic measurements, respectively. A 15 min gradient was used for separation of polar and lipophilic compounds. The mobile phases for separation of polar and semi-polar compounds were 0.1% formic acid in H_2_O (buffer A) and 0.1% formic acid in acetonitrile (buffer B). The chromatographic separation of these analytes was performed in the following conditions: A 99% initial to 1min, A 99% to A 60% to 11 min, A 60% to A 30% to 13 min and A30% to A 1% to 15 min. The following mobile phases were used for lipids and lipophilic metabolites separation: 1% of 1M NH4Ac in 0.1% acetic acid (buffer A) and acetonitrile: isopropanol (7:3) containing 1% of 1M NH4Ac in 0.1% acetic acid (buffer B). The separation of lipids and lipophilic compounds was performed with a step gradient from 45% initial to 1min, 45% A to 25 % A in 4 min, 25% to 11% A in 11 min and 11% to 0% A in 15 min. Chromatograms were recorded in full scan MS mode (Mass Range [100-1500]).

All mass spectra were acquired in positive and negative mode with the following settings of the instrument: heated electrospray ionization (HESI) was used, spray voltage was 3.5 kV, capillary temperature 275 °C, sheath gas flow rate 60 units, mass resolving power 70000, 3e6 target value (AGC) and maximal fill time of 200 ms. After measurement, samples were pooled for lipid annotation. The fragmentation was performed in data dependent MS/MS mode using higher-energy collisional dissociation (HCD). The three most intense ions were selected for fragmentation per cycle using normalized collision energy of 25 eV. The full scan spectra were acquired in the 300-1500 m/z range at a mass resolution of 35000 with an AGC of 1e5 ions with maximal fill time of 100 ms. The target values for data dependent MS/MS scans were set to 5e4 ion with a maximal fill time of 50 ms and an isolation window of 1. The MS/MS spectrum were recorded in the mass range from 50 to 1500 m/z. The MS/MS ions were measured at a resolution 17500 and the dynamic exclusion was set to 3s. Thermo Excalibur was used for the data acquisition.

#### 2.6.3. LC-MS metabolite data pre-processing

Extraction of the peak lists with mass-to-charge ratio and retention time from the LC-MS raw data files was performed using REFINER MS® 11.1 (GeneData, Basel, Switzerland) with the following workflow: Data Sweep, Grid, Chromatogram Noise Subtraction, Intensity thresholding, Chromatogram RT alignment and Chromatogram Peak Detection. An in-house software was used to align and filter the data.

#### 2.6.4. LC-MS metabolite data annotation

An in-house metaSysX database of chemical compounds was used to match the features detected in the LC-MS polar and non-polar platform. The metaSysX database contains the mass-to-charge ratio and the retention time information of reference compounds measured at the same chromatographic and spectrometric condition as sample measurements. A 5 ppm and 0.06 min deviation from the reference compounds mass-to-charge-ratio and retention time respectively were used as matching criteria for polar platform and 5 ppm and the time deviation of 0.055 for lipid annotation. Co-eluting compounds with the same ion mass were kept as unsolved annotations. Lipid annotation was additionally confirmed by MS/MS generated fragments using a metaSysX-developed R-based algorithm. The algorithm uses information from fragments, neutral losses, presence and precursor masses detected in both ionization modes. This information was combined with the information obtained from the database search to give an additional level of robustness to the data. The lipid database was created based on the precursor ion mass, fragmentation spectrum and elution patterns (Bromke et al., 2015). The elution patterns of annotated lipids were used to provide an additional level of robustness to the lipid annotation.

#### 2.6.5. LC-MS metabolite data analysis

Metabolite peaks were normalized on the basis of sample mean by: log_2_(intensity)-median(log_2_(intensity of the whole sample) + median(log_2_(intensity of all samples in the structure). Samples were normalized to the sample median per organism by: log_2_(intensity of metabolite)-median(log_2_(intensity of the sample)) + median(log_2_(intensities of all metabolites in the organism)). Since all experiments were performed using four samples, missing peaks were handled as follows: when one peak was missing from a sample, it was estimated as the median of the remaining three. If a peak was missing from more than one sample, they were handled as missing values. When peaks were missing from all four samples, they were all set to zero (0). Per metabolite, we performed a two-sample *t*-test assuming unequal variance on the log-transformed peak heights and we applied an FDR-correction at 5% to the p-values.

### 2.7. Quantification of defence-associated hormones and marker gene transcripts in co-infested leaves

To assess if *T. urticae*-induced defence responses on bean were suppressed by *T. evansi* feeding, a phenomenon observed on tomato (Alba et al., 2015), we carried out co-infestation assays where both mite species co-occur on the same leaf, using two approaches that differed in the spatial scale of mite infestation. First, we created five treatments of mite infestation: 1) leaves infested with either 30 *T. evansi* (Te); 2) with 30 *T. urticae* (Tu); 3) with 30 mites of each species, without a lanolin barrier that separates them (CoMix); 4) with 30 mites of each species, separated from each other by a lanolin-glue mixture (1:1, v/v) perpendicular to the midrib, with *T. evansi* residing at the basal section (CoTe) and *T. urticae* at the tip section (CoTu) of the leaf; and 5) whole leaves of un-infested plants as controls. Regardless of the treatments, all leaves received a lanolin-glue barrier around the petiole. Assays were conducted in two experimental blocks with a total of 9-10 plants per treatment. For the second co-infestation approach, mites were confined to a much smaller leaf area. That is, the lanolin-glue mixture (1:1, v/v) was used to create a single circular arena (Ø 20 mm) per leaf, centred on the adaxial side and crossing the midrib. Each arena was then either infested with 1) 10 *T. evansi* (Te); or with 2) 10 *T. urticae* (Tu); or with 3) 10 mites of each species, without any restrictions (CoMix), or where 4) left un-infested as controls. This experiment was performed in three blocks, with and 15-21 plants were used per treatment in total. For both experimental approaches, two days after plants were infested, leaves or leaf sections (depending on the approach) were harvested, flash-frozen in liquid nitrogen and stored at -80°C until further analyses.

Phytohormones were isolated from approximately 200 mg leaf tissue and analysed by means of liquid chromatography-tandem mass spectrometry (LC-MS/MS) according to the procedures described by Schimmel *et al*. (2017a), with the exception that the residue was resuspended in 0.5 ml 70% methanol (v/v). Data from the two experimental blocks were analysed together and log-transformed before analysis. Leaf material for gene expression and phytohormone analysis were collected independently. Total RNA was isolated from approximately 200 mg bean leaf tissue using the hot phenol method (Verwoerd et al., 1989). DNAse treatment, cDNA synthesis and quantitative polymerase chain reactions (qPCRs) were performed according to the procedures described by Schimmel *et al*. (2017a).

Using the RNAseq data generated in this study (Section 2.5), we selected two SA- and two JA-responsive marker genes for quantification of expression using qPCR. To select the JA markers, we searched the data for orthologs of known defence genes in other plant species and that are among the top genes (estimated by the average read counts) that respond to JA and to Tu, but not to SA. We selected one Kunitz-type proteinase inhibitor (*PI-KU;* Phvul.011G166500.1) and a terpene synthase (*TPS14;* Phvul.006G195600)) that complied with these criteria. To select the SA-markers we searched for genes that responded to SA and to Tu, but not to JA. We selected an acidic chitinase (*ChitA;* Phvul.011G167300.1), a pathogenesis-related protein (*PR1;* Phvul.006G196900.1). *Actin* (*ACT11*) was used as a reference gene to normalize data across samples. Primers used for qPCR are listed in **Table 1**. The qPCR-generated amplicons were sequenced to verify primer specificity. Differences in normalised gene expression and phytohormone concentrations were analysed using a linear model with treatment as fixed factor using the command *aov* in R (v 3.5.1).

**Table 1.**
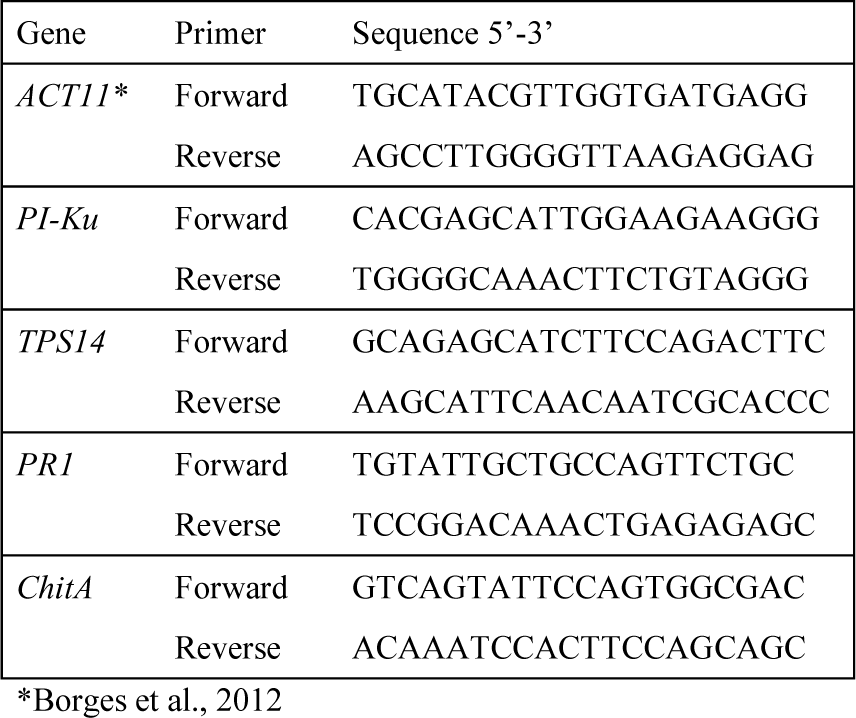
Primer sequences used for defense gene expression analysis.

### 2.9. Mite behaviour on co-infested leaves

To assess if mite behaviour is affected by the presence of competitors on the same leaf, we analysed the spatial distribution of female mites on single-species leaf infestations and in co-infested leaves. Because adult female mites are mostly feeding when they are not moving, we infer that the location of the mites reflects their preferred feeding site (Bensoussan et al., 2016). Bean leaves were infested with either 30 *T. evansi*, with 30 *T. urticae*, or with 30 mites of each species. A lanolin-glue mixture (1:1, v/v) was applied around the leaf petiole, but no further restrictions were set on mite movement across the leaf lamina. Ten plants were used per treatment (n = 10). After two days, infested leaves were excised and photographed with a Nikon D750 camera equipped with a Nikkor 60 mm f/2.8 G lens (Nikon, Tokyo, Japan), together with a scale reference. Differences in the body colouration of *T. urticae* and *T. evansi* adult females allow to easily discriminate between species visually. Thus, we measured mite position in the resulting pictures as the distance in millimetres to the petiole attachment of the leaf per individual using ImageJ (version 1.47v). The mite positions were projected on a Cartesian plane, relative to the position of the petiole, which was set to coordinates x:85, y:40. Mite positions were then expressed in coordinates and distribution heat maps were created using all the replicates per treatment using ggplot2 (Wickham, 2016) in R (v 3.5.1). Differences in the distance of mites to the petiole, in the length of the leaves used for the set up and in the number of mites alive by the end of the experiment, were analysed using a linear model with treatment as fixed factor using the command *aov*. Post hoc comparisons between treatments were conducted using the command *TukeyHSD*.

## 3. Results

### 3.1 T. urticae benefits from the presence of T. evansi on bean

The suppression of plant defences can be exploited by co-existing competitors, who can benefit from a plant with impaired defences. We investigated whether the reproductive performance of the defence-inducing mite *T. urticae* was affected when sharing a leaf with the defence-suppressing mite *T. evansi*. We found that the oviposition of *T. urticae* is significantly affected by the identity of the mite sharing their leaf (F_2, 63_ = 9.2, p < 0.001, **Fig. 1**). The number of eggs laid by *T. urticae* was ∼27% higher on leaves co-infested with *T. evansi* than on leaves co-infested with conspecifics. The oviposition rate was intermediate on control leaves, not co-infested with other mites. The percentage of surviving *T. urticae* females was not affected by the presence of other mites on the leaf (control: 76.5% ± 4.1% (mean ± SEM); co-infested with *T. urticae*: 73.0% ± 4.5%; co-infested with *T. evansi*: 75.0% ± 4.1%).

**Fig. 1.**
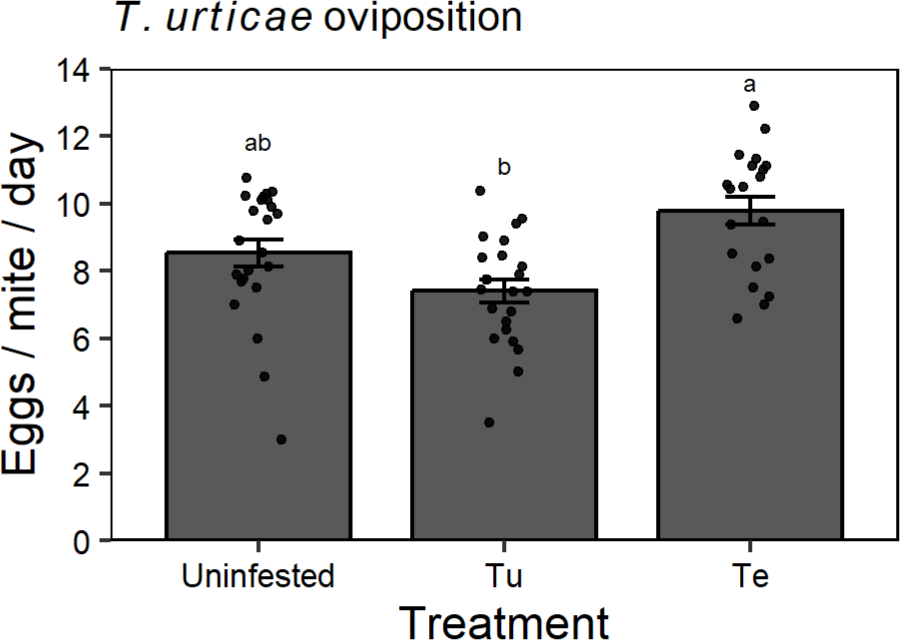
Reproductive performance of *Tetranychus urticae* on bean leaves shared with competitors. Using artificial barriers, leaves of intact bean plants were divided into a basal section and a tip section. The number of eggs laid by *T. urticae* females per day (y-axis) in the tip section was scored on leaves where the basal section was either co-infested with *T. evansi* (Te), co-infested with *T. urticae* (Tu), or left un-infested. Bars show the average number of eggs laid per *T. urticae* female per day ± standard error of the mean (SEM), with individual data points plotted over the bars. Different letters above the bars indicate significant differences between treatments according to a linear model and post hoc comparisons using a Tukey correction for multiple testing.

### 3.2 Spider mites are susceptible to the JA-dependent responses of bean

To understand the extent of mite resistance or susceptibility to bean defences, we injected leaves on intact plants with either JA or SA solutions, and then infested these plants with either *T. urticae* or *T. evansi* to assess their reproductive performance. When performing the experiment on whole leaves, the phytohormone treatment had an effect on *T. evansi* oviposition (F_2, 40_ = 5.40, p = 0.008), but not on *T. urticae* oviposition (F_2,44_ = 2.27, p = 0.11). The JA treatment reduced *T. evansi* oviposition by ∼32% compared to the mock treatment, but the SA treatment did not affect *T. evansi* oviposition (**Fig. 2a**). When performing the experiment on leaf discs, there was a significant effect of the phytohormone treatments on both *T. evansi* (F_3, 309_ = 46.87, p < 0.001) and on *T. urticae* oviposition (F_3, 332.04_ = 3.06, p = 0.03). The JA treatment reduced *T. evansi* oviposition by ∼26% and the SA+JA treatment by ∼18% compared to the mock treatment, but SA had no effect. Only the JA treatment reduced *T. urticae* oviposition by 13% compared to the mock treatment, and there was no significant effect of the SA or the SA+JA treatments (**Fig. 2b**). Overall, we found that mite performance is negatively affected upon bean treatment with JA, but not with SA, independently of the mite species. We also compared the proportion of missing mites, i.e. the mites that died, got stuck, or went missing during the experiment. We found that ∼8% more *T. urticae* females were alive by the end of the experiment on whole leaves for the SA treatment compared to the control (χ^2^ = 5.4936, df = 1, p = 0.019; **Fig. 2c**).

**Fig. 2.**
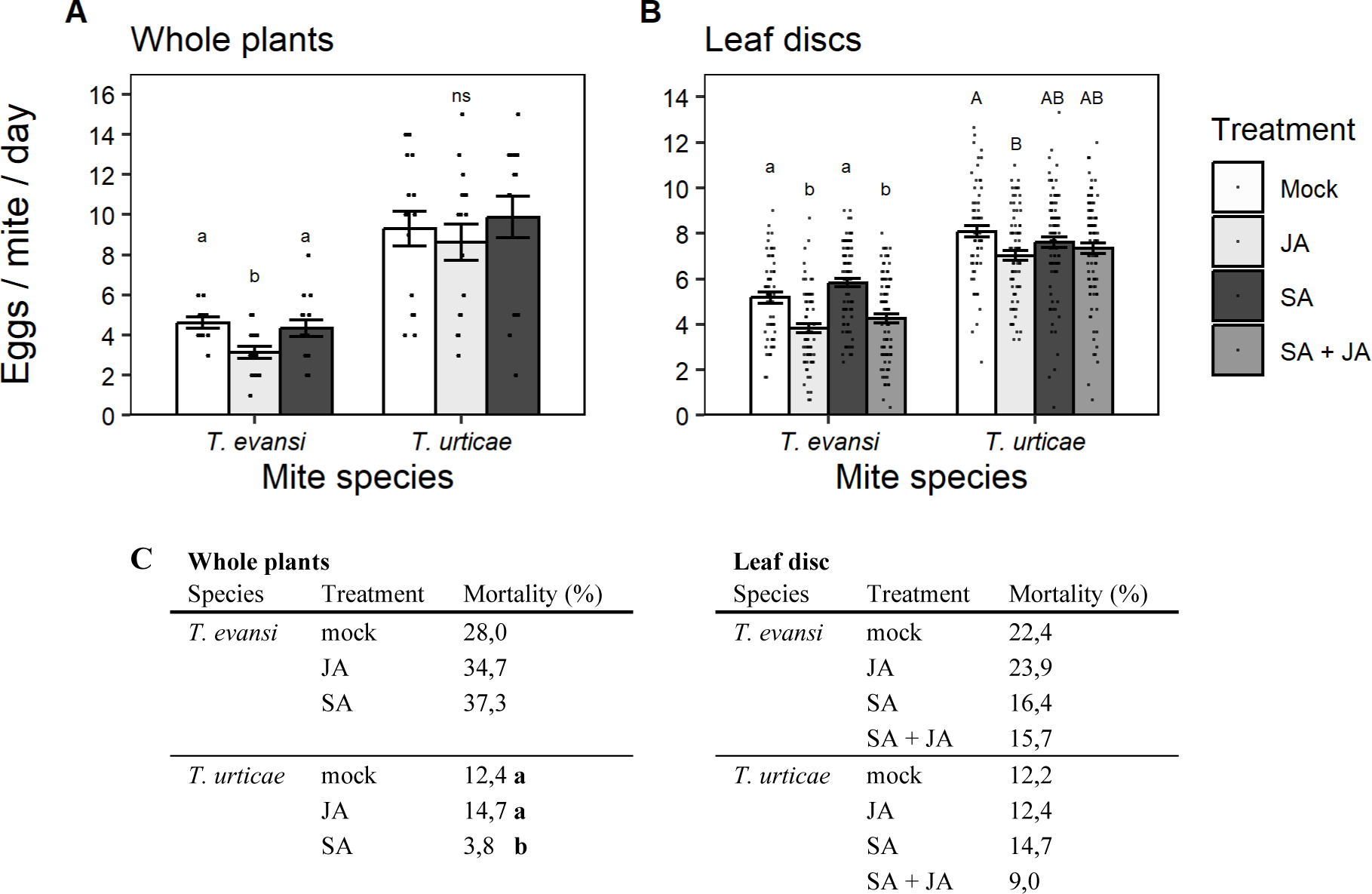
Reproductive performance and survival of adult female mites on bean upon exogenous phytohormone application. A) The number of eggs laid per day (y-axis) by *T. evansi* (Te) and *T. urticae* (Tu) females was scored on whole bean plants treated with jasmonic acid (JA), salicylic acid (SA), a combination of both (SA+JA), or a control solution (Mock; legend). B) The number of eggs per mite per day of both species tested on cut out leaf discs from phytohormone-treated bean plants (legend). Different letters above bars indicate significant differences according to a linear mixed-effects model followed by a Tukey correction for multiple comparisons per mite species. ns, not significant. C) The mortality of female mites from both species during the experiments. Different letters next to percentages indicate significant differences within species due to treatment based on a chi-square test.

### 3.3. Molecular changes on bean upon phytohormone application and mite infestation

To characterise the molecular responses of bean to mite feeding, and the role of JA and SA in these responses, we conducted transcriptomic and metabolomic analyses on bean plants treated either with phytohormones or infested with mites.

#### 3.3.1. Transcriptional changes

We collected RNA from leaves 12 hours after infiltration with JA or SA, or 4 days after infestation with *T. urticae* or *T. evansi*, using un-infested and un-infiltrated plants as controls. A total of 20,371 transcripts passed our selection criteria and were used for the analysis. We analysed these transcripts as genes that responded only to phytohormones, genes that responded to both mite feeding and phytohormones, and mite-responsive genes. The genes that were significantly different from controls are further analysed (**Fig. 3A-B**). Results included in tables 2-13 exclude genes without annotation.

**Fig. 3.**
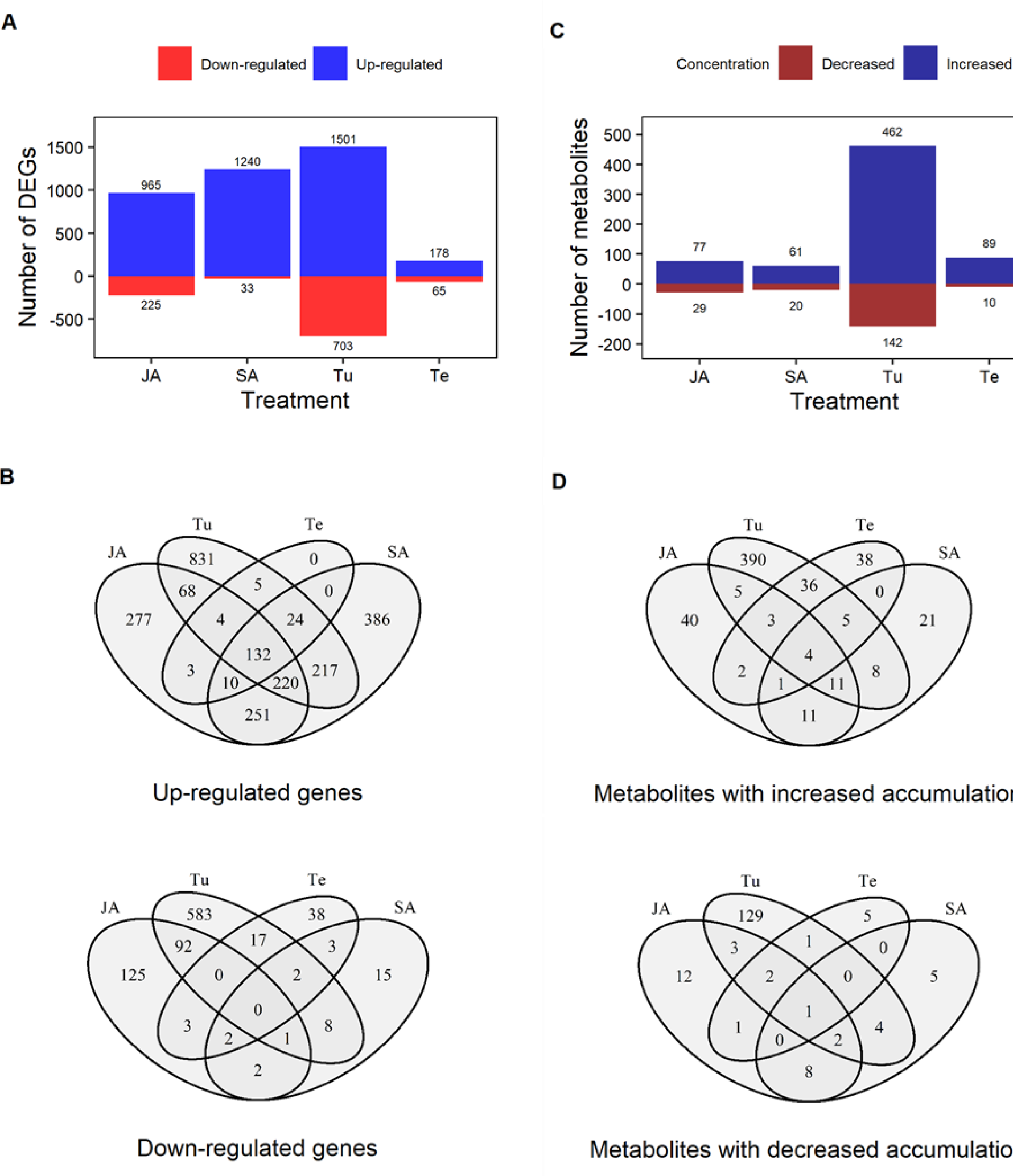
Transcriptional and metabolic changes in bean plants exposed either to phytohormones or mite feeding. Bean leaves were treated for 12 hours with the phytohormones jasmonic acid (JA), salicylic acid (SA), or were infested with either *T. urticae* (Tu) or *T. evansi* (Te) for four days. Un-infested, un-treated plants served as controls and were used as a common reference in the comparisons. A) Total number of genes with a significantly different expression compared to controls (y-axis) across treatments (x-axis). Numbers of differently expressed genes are presented above and below the bars for up-regulated and down-regulated groups (legend). B) Venn-diagrams of genes that were up-regulated (above) and down-regulated (below) in the four treatments (JA, SA, Tu and Te) showing the overlap in transcriptomic responses of bean to the treatments. C) Total numbers of metabolites (y-axis) that accumulate to different concentrations than controls over four treatments (x-axis). Numbers of metabolites with an increased or decreased accumulation significantly different than controls are presented above and below the bars (legend). D) Venn-diagrams for the four treatments showing the overlap of the metabolic responses of bean to the treatments.

Compared to untreated controls, JA up-regulated 965 and down-regulated 235 genes, and SA up-regulated 1,240 and down-regulated 33 genes. *T. urticae* up-regulated 1,501 genes and down-regulated 703 genes, whereas *T. evansi* up-regulated 178 genes and down-regulated 65 (**Fig. 3A**). Of the JA-inducible genes, 136 were up-regulated and 2 down-regulated by both mite species. Further, 288 were uniquely up-regulated and 158 uniquely down-regulated by *T. urticae*, whereas *T. evansi* uniquely up-regulated 13 and down-regulated 7 of these genes. Of the JA-down-regulated genes only 20 were up-regulated and 93 were down-regulated by *T. urticae*. *T. evansi* did not up-regulate any of these genes and down-regulated only 8. *T. evansi* uniquely up-regulated 10 and down-regulated 7 SA-inducible genes. Of the SA-down-regulated genes only 8 were up-regulated and 9 were down-regulated by *T. urticae*. By contrast, *T. evansi* did not up-regulate these genes and down-regulated only 2 genes.

Defense-associated genes that were induced by JA in bean include subtilisin-like serine endopeptidases, lipoxygenases, and terpene synthases (**Table S1**). Among the SA-induced genes there were receptor-like kinases (RLKs), leucine rich repeats (LRRs), WRKY transcription factors, and several pathogenicity related (PR) genes (**Table S2**). Both hormones also had negative effects on expression. Among others, JA suppressed expression of multiple LRRs (**Table S3**) and SA suppressed a terpene synthase (**Table S4**). Of the genes that were up-regulated by JA or SA in bean, 39% was up-regulated by both hormones, among which a large set of chalcone and stilbene synthases, indicating the up-regulation of flavonoid biosynthesis (**Table S5**), which were also induced by *T. urticae* feeding. Among the genes that were induced by *T. urticae*, but not by *T. evansi* were RLKs, LRRs, WRKYs, and Cytochrome P450s (**Table S7**). Genes that were down-regulated by *T. urticae* were MYB and RAD-like transcription factors and also many chloroplast-located proteins (**Table S8**). Feeding by *T. evansi*, but not *T. urticae*, induced the expression of a calmodulin-binding protein, a homeobox leucine zipper, and an aquaporin-like protein (**Table S9**), while uniquely suppressing the expression of nitrate responsive Laccase 7, a phosphate transporter involved in sugar transport, a TPS and a RLK (**Table S10**). When filtering for genes induced by both mites, we observe that most of these genes are also induced by JA, SA, or both hormones and that they were induced to the same or higher level by *T. urticae* than *T. evansi*. Many of these genes have putative functions in stress, wound, and/or defence responses (**Table S11**). When filtering for genes down-regulated by both mite species, we also found multiple genes putatively involved in stress and defence responses (**Table S12**).

#### 3.3.2. Metabolomic changes

We performed an untargeted metabolomics analysis to determine if the patterns observed at the transcriptomic level were similar on the metabolomic level. Metabolites were measured for leaf samples infiltrated with JA or SA, or infested with *T. urticae* or *T. evansi*, and were collected in a similar way as the samples for the RNAseq. In the analysis, a total of 6.307 metabolites were detected.

Compared to the control, JA increased the concentration of 77 metabolites and decreased the concentration of 29 metabolites. SA increased and decreased the concentration of 61 and 20 metabolites, respectively. *T. urticae* significantly increased accumulation of 462 metabolites and decreased accumulation for 142 metabolites. *T. evansi* increased the accumulation of 89 metabolites and decreased the accumulation of 65 metabolites (**Fig. 3c**). Of these last metabolites 48 had and increased and 4 had a decreased accumulation by both species (**Fig. 3d**). We observed that JA, SA, and *T. urticae* increased the expression of genes involved in flavonoid biosynthesis (see section 3.3 *Transcriptomic changes*), so we searched for changes in metabolites associated with the accumulation of flavonoids in our samples. We found 14 flavonoids that exclusively accumulate to higher concentrations upon infestation with *T. urticae* than controls or treatments with *T. evansi* (**Table 3**).

**Table 3.** The name, class, and average peak area of metabolites associated with flavonoid biosynthesis, accumulated only in the presence of *T. urticae*—not accumulated under control conditions or by *T. evansi*.

| Metabolite | Class II | Treatment* |  |  |  |  |
| --- | --- | --- | --- | --- | --- | --- |
|  |  | Control | SA | JA | Tu | Te |
| Amentoflavone | Biflavonoids and polyflavonoids | ND | ND | ND | 175.526<sub>4</sub> | ND |
| Apigeninidin | Anthocyanidins and anthocyanins | ND | ND | ND | 576.493<sub>4</sub> | 156.757<sub>1</sub> |
| Phloretin | Dihydrochalcones | ND | ND | ND | 122.174<sub>2</sub> | ND |
| Apigenin 7-O-beta-D-glucoside | Flavones | ND | 108.546<sub>1</sub> | ND | 130.006<sub>4</sub> | ND |
| Isovitexin | Flavones | ND | 304.565<sub>4</sub> | 268.890<sub>4</sub> | 250.777<sub>4</sub> | 202.323<sub>1</sub> |
| Luteolin 7-O-beta-D-glucoside | Flavones | ND | ND | ND | 216.709<sub>4</sub> | ND |
| Fisetin | Flavonols | ND | ND | ND | 81.748<sub>2</sub> | ND |
| Coumestrol | Coumestanes | ND | ND | ND | 8.035.416<sub>4</sub> | 315.944<sub>1</sub> |
| Calycosin | Isoflavones | ND | ND | ND | 2.965.654<sub>4</sub> | ND |
| Daidzein | Isoflavones | ND | ND | ND | 4.526.690<sub>2</sub> | ND |
| Genistein 7-O-beta-D-glucoside | Isoflavones | ND | 2.712.748<sub>4</sub> | 1.820.395<sub>4</sub> | 1.759.725<sub>4</sub> | 429.102<sub>1</sub> |
| Glycitein | Isoflavones | ND | ND | ND | 71.192<sub>3</sub> | ND |
| Glycitin | Isoflavones | ND | ND | ND | 265.916<sub>4</sub> | ND |
| Ononin | Isoflavones | ND | ND | ND | 387.734<sub>4</sub> | ND |

\*‘ND’ means not detected; subscript indicates the number of replicates in which metabolites were detected. Metabolites that were detected in at least 2 of the 4 replicates are highlighted in black.

### 3.4. Defence responses of bean upon single-species infestations and two-species co-infestations

#### 3.4.1. Phytohormone accumulation

To investigate the ability of *T. evansi* to manipulate bean defences, we measured the concentrations of JA, JA-Ile, 12-oxo-phytodienoic acid (OPDA) and SA under single *T. evansi* (Te) or *T. urticae* (Tu) infestations, and under co-infestation conditions, with either both mites sharing a whole leaf (CoMix), or with *T. evansi* and *T. urticae* separated on different areas of the same leaf (CoTe and CoTu, respectively; **Fig. 4**). Overall, JA-related phytohormones did not differ significantly between the control, the Tu and the CoMix treatments; a similar trend was observed with Te feeding, except only OPDA was significantly lower than control (**Fig. 4A-C**). The CoMix treatment, where both mites species fed freely from the leaves, did not differ from Tu feeding for any phytohormone, but it accumulated significantly higher concentrations of OPDA and SA than Te feeding. The concentration of SA increased upon infestation with both single-species treatments compared to controls, but Tu and CoMix resulted in SA concentrations over four times higher than Te (**Fig. 4D**). JA-related hormones were accumulated to similar levels between the CoMix treatment and the two CoTe and CoTu treatments. While SA concentrations were similar between the CoMix and the CoTu treatments, SA concentrations were lower on the CoTe than CoMix and CoTu.

**Fig. 4.**
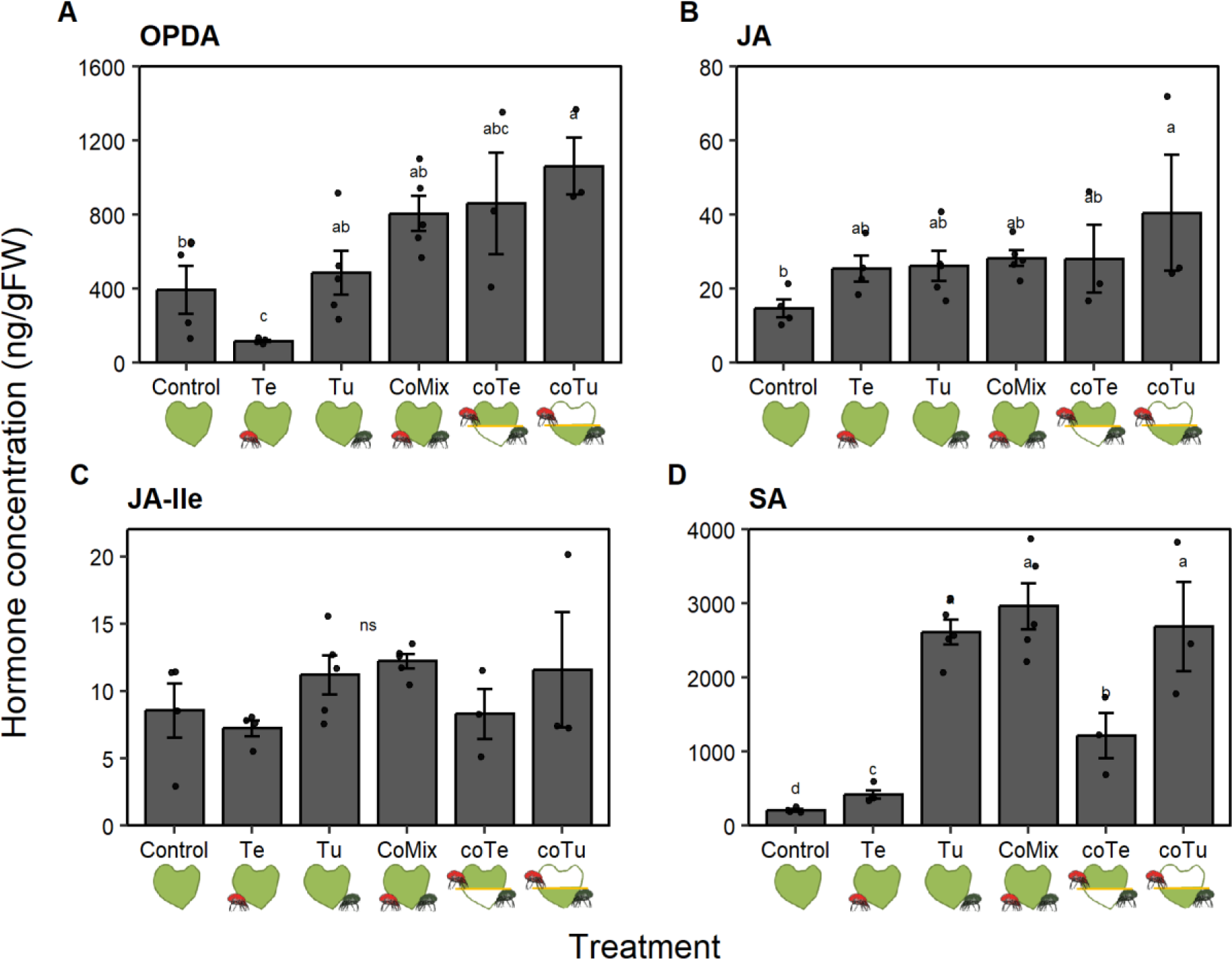
Phytohormone concentrations on bean measured after treatments with single-species or two-species mite infestations. Concentration (ng/g fresh weight FW; y-axes) of A) 12-oxo-phytodienoic acid (OPDA), B) jasmonic acid (JA), C) JA-isoleucine (JA-Ile), and D) salicylic acid (SA) in treated bean plants that were either left un-infested (Control), infested with *T. evansi* (Te), *T. urticae* (Tu), with 30 *T. evansi* and 30 *T. urticae* without restrictions on the leaf (CoMix), or with *T. evansi* infesting the basal half of the leaf (CoTe) and *T. urticae* infesting the tip section of the leaf (CoTu; x-axis). Leaf tissue of CoTe and CoTu were collected separately. Different letters above the bars within each panel indicate significant differences of the log-transformed data according to a linear mixed-effects model followed by a Tukey correction fo multiple comparisons. n=3-5

#### 3.4.2. Quantification of defence marker gene expression at two different spatial scales

From the transcriptomic data, we selected genes that could be used as defence markers for qPCR quantification. We selected two candidates that responded to JA application and to Tu feeding, but not to SA application. From the 11 loci in annotated as Kunitz-type proteinase inhibitors (*PI-KU*), four complied to our selection criteria. We designed primers for the two genes with the highest average read counts (Phvul.004G129900.1 with 5071 reads and Phvul.011G166500.1 with 863 reads). Since only the primers for the second loci resulted in a successful measurement, we continued with this locus. As a second marker gene we selected a terpene synthase. 15 loci are annotated as a terpene synthase on bean, out of which Phvul.002G219300.1 was represented by the highest number of average reads (915), but this locus also responded marginally to SA. The second locus (Phvul.006G195600.1 [*TPS14*] with 617 reads on average) did not respond to SA and thus was chosen as a second JA marker (*TPS14*). For the SA-markers, we found 21 chitinases that responded to exogenous SA application, and selected Phvul.011G167300.1 (an acidic chitinase, *ChitA*) since the 5 loci with higher read counts (all above 11330 as for this locus) all were basic chitinases that also responded to JA. We also selected a pathogenesis-related protein (*PR1*) locus as a SA marker. We found 7 *PR1* loci in the data set. Locus Phvul.006G197200.1 and Phvul.006G196900.1 had the highest read counts (146940 and 84511, respectively), and both responded to SA but not to JA application. Since only the primers for the second locus amplified a fragment, we continued with the locus (*PR1*). Note that after FDR correction for multiple comparison, *TPS14* was significantly expressed for JA and Tu; the *ChitA* and the *PI-KU* only for Tu; and the *PR1* for neither. Yet, since they were all significant for the relevant treatments, we considered the absolute read count as an important criterion for reliable detection (**Table S13**).

To further our insight into the extent of induction and suppression of bean defences by mite feeding and the spatial extent of this effect, we measured the expression of defence marker genes by single-species and co-infestation treatments both on whole plants and on leaf discs. There was a significant effect of mite treatment for the four marker genes both in whole leaves and leaf discs (**Table S14**). Clear patterns were observed between the single-species infestation treatments. When infesting whole leaves, Tu induced the expression of all JA and SA marker genes compared to controls, although only marginally for *TPS14* (**Fig. 5A**), whereas Te did not induce any of these genes (**Fig. 5**). The treatment with both species co-infesting a leaf without any barriers (CoMix) showed a significantly higher expression of all genes compared to controls and to Te, while the gene expression of the CoMix did not differ from Tu for most genes, with the exception of *PR1*, where the expression was higher in CoMix than in Tu, Te, and control (**Fig. 5C**). Although the expression of *TPS14*, *PI-KU*, and *ChitA* in CoMix did not differ significantly compared to Tu, it showed a trend towards higher expression for *TPS14* and towards lower expression for *PI-KU* and *ChitA*. In co-infestation treatments where mite species were isolated to different areas of the leaves via a lanolin barrier, *T. evansi* occupied the basal half of the leaf (CoTe) and *T. urticae* infested the tip of the leaf (CoTu). We measured gene expression on the tissue of leaf where only one of the two species was present. Leaf halves with *T. urticae* (CoTu) showed a level of expression comparable to leaves co-infested with mites of both species (CoMix), although the expression of *PI-KU* and *ChitA* was significantly higher in the CoTu part of the leaf than in CoMix treatments (**Fig. 5B, 5D**).

**Fig. 5.**
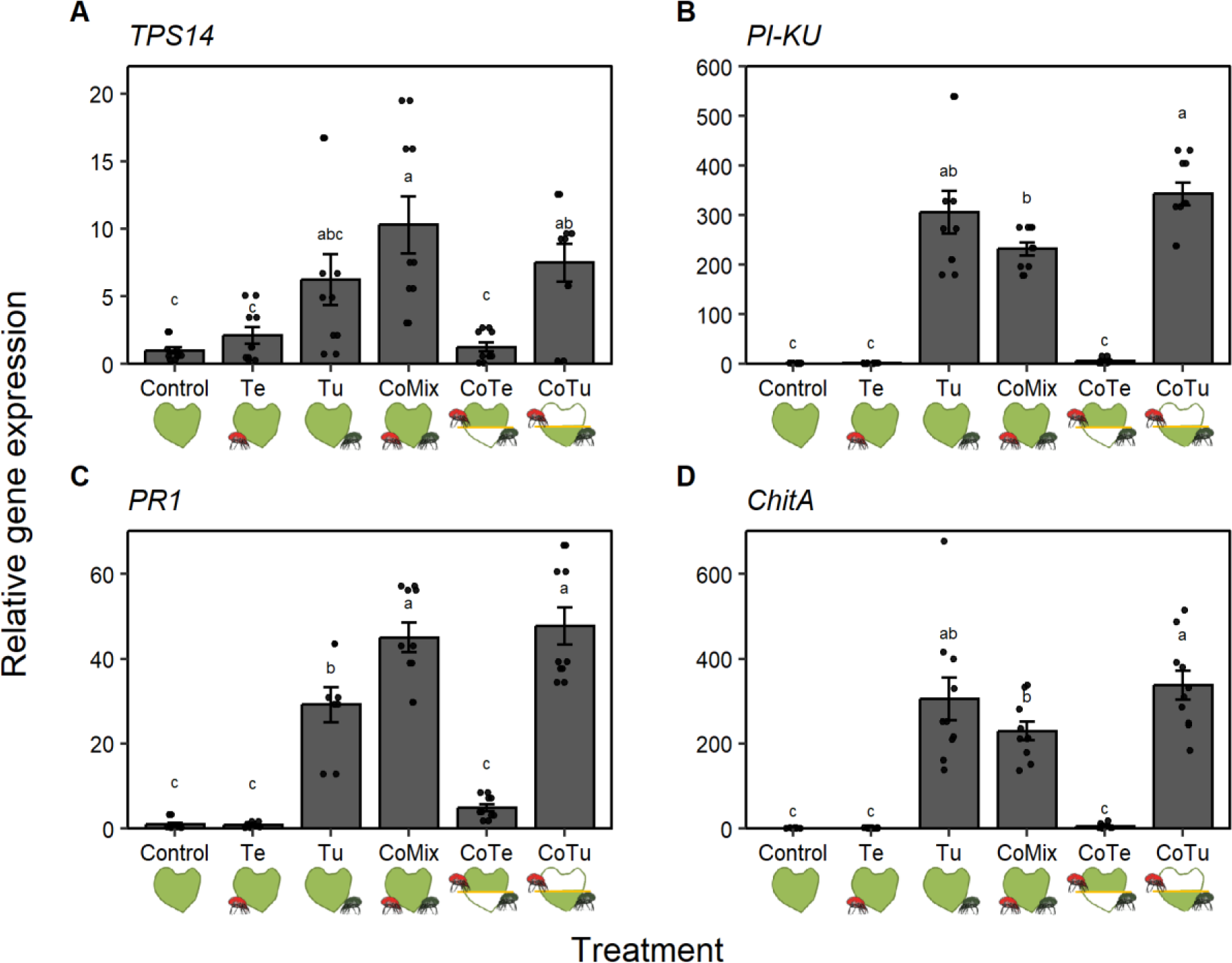
Defence marker expression on bean measured after treatments with single-species or two-species mite infestations. Expression of defense marker genes relative to *actin* (y-axes) of JA-responsive markers A) *terpene synthase 14* (*TPS14*) and B) *Kunitz trypsin inhibitor* (*PI-KU*), and SA-responsive markers C) *pathogenesis-related protein 1* (*PR1*) and D) *chitinase A* (*ChitA*) across treatments (x-axes): un-infested leaves (Control), leaves infested with *T. evansi* (Te), *T. urticae* (Tu), both species without barrier to separate them (CoMix), or infested with both species but isolating them with a barrier to the basal side of the leaf for *T. evansi* (CoTe) or the tip section of the leaf for *T. urticae* (CoTu). Different letters above the bars indicate significant differences within panels of the log transformed data across three experimental blocks, according to a linear mixed-effects model followed by a Tukey correction for multiple comparisons.

To understand whether the induction and suppression of marker gene expression was locally restricted to the area closest to mite feeding, we conducted a similar experiment, but instead of allowing the mites to roam through the whole leaf, we restricted them to a small circular arena in the centre of a leaf, thereby bringing them closer together. As on whole leaves, we observed a significantly higher expression of all marker genes upon feeding by Tu, whereas gene expression by Te was not different from controls for most genes except for *PR1*, which had a higher expression than controls for Te, although to a considerably lower magnitude than Te (**Fig. 6**). At this restricted spatial scale, we also observed that CoMix treatments induced a higher expression for all genes compared to control and to Te. However, gene expression was significantly reduced in the CoMix treatment compared to Tu for *PI-KU* and marginally for *ChitA*, and a similar non-significant trend was observed for *TPS14* and *PR1*, despite the fact that CoMix treatments had double the number of mites per arena than single-species treatments (**Fig. 6**).

**Fig. 6.**
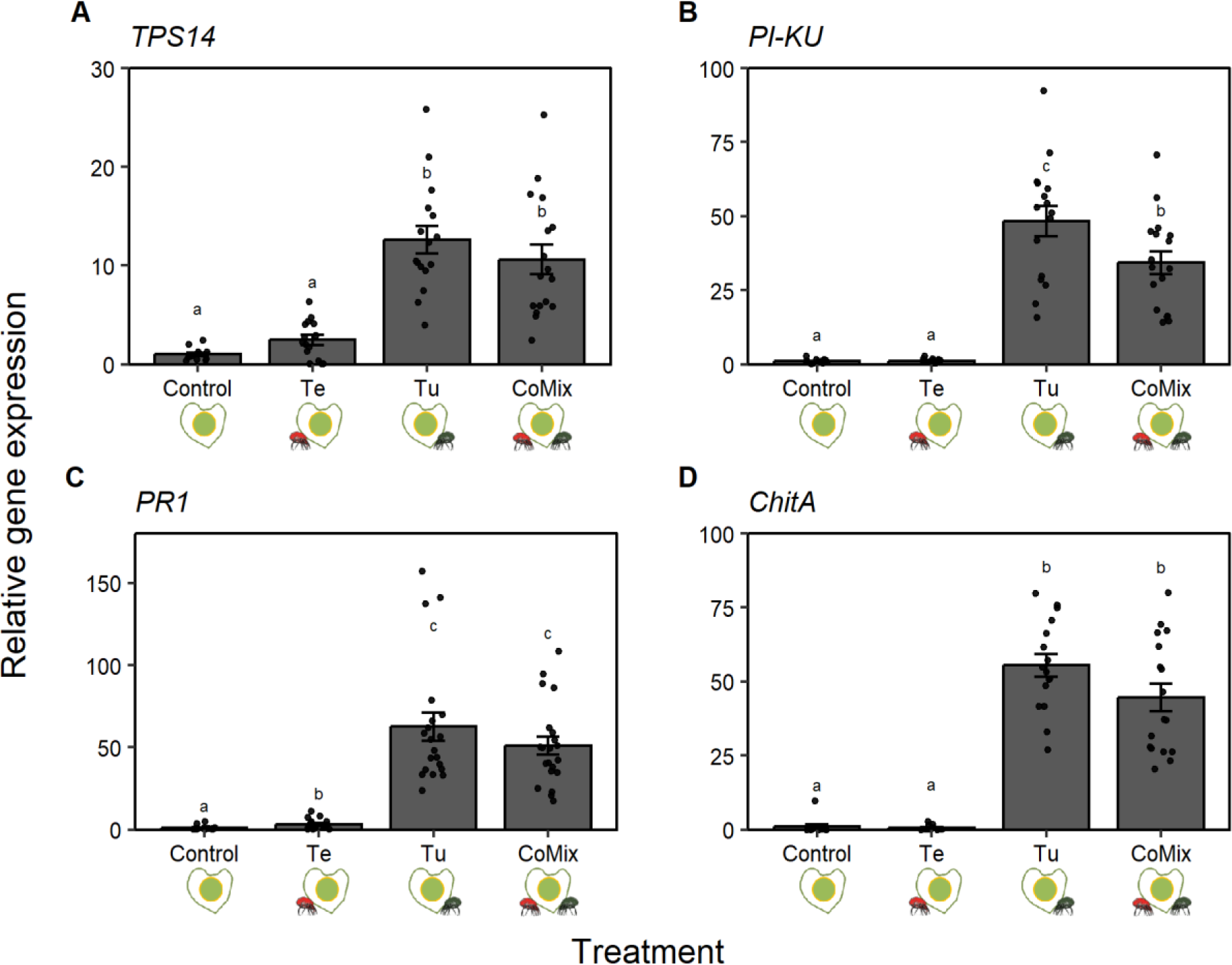
Defence marker expression on small bean arenas with single-species or two-species mite infestations. Expression of defence marker genes relative to *actin* (y-axes) of JA-responsive markers A) *terpene synthase 14* (*TPS14*) and B) *Kunitz trypsin inhibitor* (*PI-KU*), and SA-responsive markers C) *pathogenesis-related protein 1* (*PR1*) and D) *chitinase A* (*ChitA*) across treatments (x-axes): un-infested arenas (Control), arenas infested with *T. evansi* (Te), *T. urticae* (Tu), or arenas infested with both species (CoMix). Different letters above the bars indicate significant differences within panels of the log transformed data across three experimental blocks, according to a linear mixed-effects model followed by a Tukey correction for multiple comparisons (n= 12-21).

### 3.5. T. evansi displaces T. urticae from its preferred feeding site when co-infesting a bean leaf

To assess whether the presence of competitors on the same bean leaf elicits behavioural changes of each of the mite species, we scored the location on the leaves where each of the mite species fed in single-species infestations, and compared these locations to leafs co-infested with both species. We pooled the positions of each individual mite from ten replicate pictures (for individual replicates see **Fig. S1**), and found that both species have a clear preferred feeding site (**Fig. 7**). Both species tend to feed at the base of the leaf, close where the midrib connects with the petiole (**Fig. 7I and 7III**). When the two species are together on a leaf, *T. evansi* individuals maintain a similar spatial distribution as when they are alone on a leaf (**Fig. 7I-II**). However, in co-infested leaves *T. urticae* moved away from its preferred feeding site close to the base of the leaf, further towards the leaf laminae and away from the site where *T. evansi* fed (**Fig. 7III-IV**). We used the data of individual mite positions to compare the average distance to the petiole for each mite species, either in single-species infestations or in co-infestations (**Fig. 8**). We found that *T. evansi* individuals clump close to each other and feed closer to the petiole than *T. urticae* does in single infestations (**Fig. 8A-B**). During co-infestations, *T. evansi* maintains its preferred feeding site relative the petiole (Fig. 8), whereas *T. urticae* moves away from the area close to the petiole, into the leaf lamina and away from the feeding site occupied by *T. evansi* (**Fig. 7II-IV, Fig. 8A-B**), causing a shift in the distribution of *T. urticae* mites in co-infested leaves compared to leaves infested only with *T. urticae*. There were no significant differences in average leaf length or average number of mites between the treatments (**Fig. 8C-D**).

**Fig. 7.**
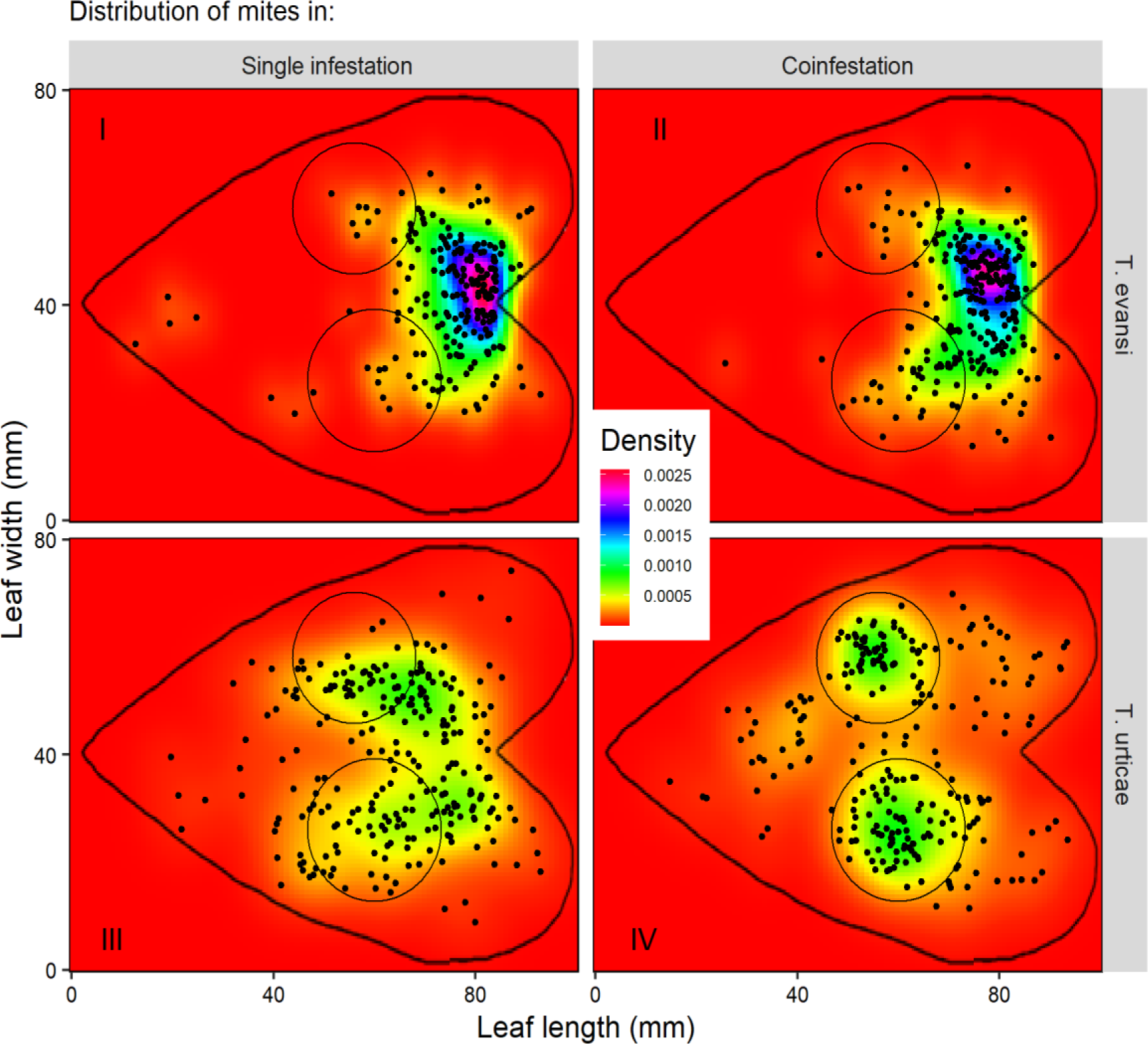
Spatial distribution of individual mites in single-species infestations and in co-infestations with competitors. Heat maps compiling the distribution of *T. evansi* (panels I and II) and *T. urticae* (panels III and IV) individuals 48 hours after infestation, when feeding alone on a leaf (panels I and III) or in co-infestations with *T. urticae* and *T. evansi* together (panels II and IV) on the abaxial side of a bean leaf. The tilted heart shape represents a hypothetical leaf shape, and the two circles represent the places with the highest densities of *T. urticae* mites when feeding with *T. evansi* (panel IV). Every dot is the position of an individual mite along the length and width in millimeters of replicate bean leaves (x-and y-axes). Positions of individuals are projected into a cartesian plane and are relative to the petiole, which is set at coordinates x: 85; y: 40. The density of individuals across the plane is represented by different colours along the gradient shown in the legend.

**Fig. 8.**
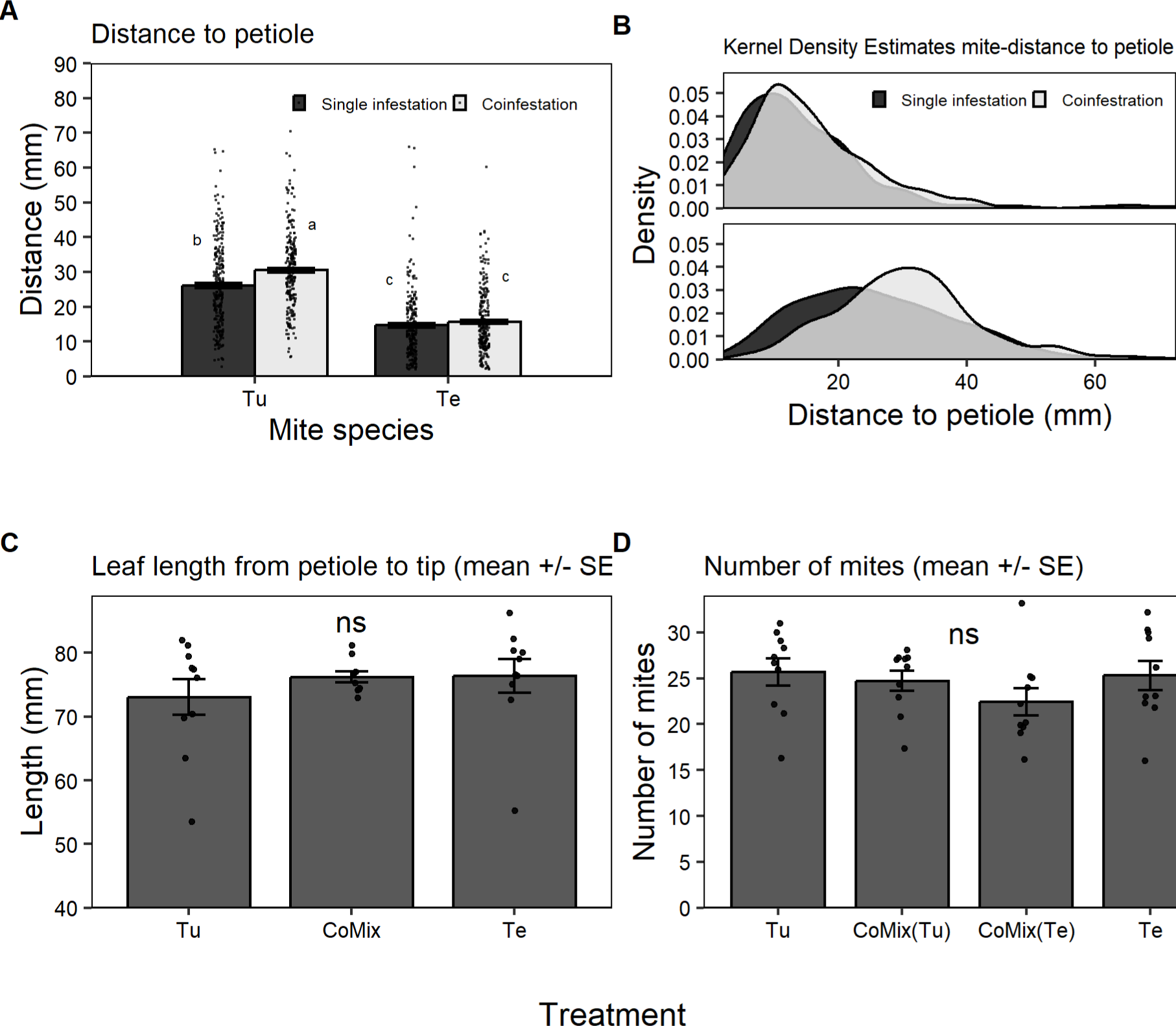
Positions of individual mites relative to the petiole of bean leaves infested either with single-species or co-infested with competitors. A) Distance from the petiole (y-axis) of *T. urticae* (Tu) and *T. evansi* (Te) individuals in single-species infestations and in co-infestations with both mite species (legend). Bars represent the average distance to the petiole in millimeters ± SEM. Individual mites are represented by dots plotted on top of the bars. Different letters above the bars indicate significant differences between treatments, using log transformed data across ten pooled experimental blocks according to a linear mixed-effects model followed by a Tukey correction for multiple comparisons. B) Kernel density estimates (y-axis) of the distribution of each mite species in single-species and co-infestation treatments for *T. evansi* (top) and for *T. urticae* (bottom) individuals plotted against the distance to the petiole (x-axis). C) Average leaf length (±SEM) measured from petiole to tip across treatments. D) Average number of mites (±SEM) that were retrieved alive at the end of the experiment across treatments. ‘ns’ = not significant.

## 4. Discussion

The suppression of plant immunity by herbivores, a strategy to cope with host defences, can have consequences that reverberate across the ecological community. For example, plant tissue impaired in defences can be open for exploitation by organisms that could not access the defended resource before, and thus suppression can alter the interactions between co-occurring competitors. The tomato specialist spider mite *T. evansi* can suppress its host defences, which in turn can facilitate tomato colonization of a non-adapted competitor mite, *T. urticae* (R. A. Sarmento et al., 2011). The host range of *T. evansi* includes species outside the family Solanaceae, but there is little evidence on whether it can suppress the defences of hosts beyond tomato, and how this suppression can alter interactions with competitors. In this study, we describe the molecular changes in bean transcripts and metabolites upon mite infestation, and study the spatial distribution of co-occurring mites that either induce or suppress defences on bean leaves. We found that I) *T. urticae* oviposition is improved on bean leaves co-infested with *T. evansi* (**Fig. 1**), II) JA negatively affects oviposition of spider mites on bean (**Fig. 2**), III) *T. evansi*, compared to *T. urticae*, has an attenuated effect on the plant in terms of transcription or metabolite accumulation (**Fig. 3**), IV) *T. evansi* can suppress gene expression of defence marker genes in bean (**Fig. 6**), and V) *T. evansi* displaces *T. urticae* from its preferred feeding site on bean leaves (**Fig. 7**).

### 4.1 Mite oviposition increases with defence suppression

In this study we have shown that the presence of *T. urticae* on the same leaf negatively affects oviposition of co-occurring conspecific mites, whereas the simultaneous presence of *T. evansi* benefits *T. urticae*’s oviposition on bean (**Fig. 1**). The results in **Fig. 4, 5, and 6** show that feeding from bean by *T. urticae* induces host defense responses and **Fig. 3** shows that the mites are at least susceptible to JA induced defenses in bean. The results also suggest that *T. evansi* is able to suppress the induced defenses of bean (**Fig. 6**). This effect has been observed before in tomato, when the two mite species were introduced simultaneously (Alba et al., 2015) or sequentially (Renato A. Sarmento et al., 2011b). Observing the improved oviposition of *T. urticae* also on bean infested with *T. evansi* suggests that suppression is mediated through the manipulation of a conserved defense mechanism, that is shared in at least Fabaceae and Solanaceae.

### 4.2 Oviposition is negatively affected by JA induced defenses

We showed that *T. urticae* induced JA and SA marker gene expression and *T. evansi* does not, or much less so than *T. urticae* (**Fig. 5**, **Fig. 6**). Bean plants treated with JA had a negative effect on the oviposition of both mite species, but plants treated with SA did not (**Fig. 2**). In tomato, however, spider mites have been shown to be negatively affected by both JA-(Ament et al., 2004) and SA-dependent defenses (Villarroel et al., 2016). From the RNAseq results we know that the JA and SA treatments clearly affect the total DEGs in the plants, with a large overlap with mite induce genes (**Fig. 3**). Overall, similarly to tomato, this shows that *T. urticae* induces plant defences and *T. evansi* does not, or at least to a lesser extent. But it shows also that *T. evansi* is susceptible to the bean defences.

### 4.3. T. evansi dampens bean response

The bean transcriptional response to the phytohormones or the spider mites was characterized mainly by up-regulation of genes (**Fig. 3**), similar to pepper (Zhang et al., 2020). In tomato, a similar number of genes were up-or down-regulated (Schimmel et al., 2018). Since mite feeding causes physical damage to a leaf, and hormones were infiltrated with a syringe, all bean transcript data (except those from controls) presented here probably also include a residual wound response. In our dataset, The *T. urticae* treatment had the most differentially expressed genes. As observed before in tomato (Schimmel et al., 2018), most of the genes up- or down-regulated in response to *T. urticae*, were not differentially regulated in response to *T. evansi*. To compare the two mites, *T. urticae* increased the expression of 1,501 genes and *T. evansi* only 178 genes and out of these only 13 are unique to *T. evansi* (**Table S9**). Ten of these 13 are responsive to both JA and SA and only three are uniquely JA responsive (**Fig. 3**). Six of these genes have an unknown function and four have a role in stress/defense response, including glutaminyl cyclase (**Table S9**) which is also induced in response to cabbage leaf curl virus (Ascencio-Ibáñez et al., 2008). The fact that the plant responds so drastically different to the two mite species suggests that *T. evansi* does not simply manipulate a specific pathway but most likely targets central regulators in the cell, as hypothesized earlier by Schimmel *et al*. (2018).

### 4.4. T. evansi suppression of gene expression

In the experiment where the distance between the two mite species was relatively large, we observed that *T. evansi* promoted *T. urticae* oviposition (**Fig. 1**) despite the fact that we did not observe suppression of defense-genes on the leaf as a whole (**Fig. 5**). What appears to be case is that the suppression of defenses occurs at different degrees on different scales. On co-infested arenas where the mites share a smaller leaf area we observe a more consistent trend towards suppression of defense marker gene expression, compared to co-infested whole leaves (**Fig. 6** vs **Fig. 5**). This effect was the clearest for the Kunitz-type proteinase inhibitor (*PI-KU*) while for the terpene synthase (*TPS14*) and *PR-1* the trends were opposite whereas ChitA was not affected by the different set-ups. This suggests that different signaling events with a different spatio-temporal organization may account for the down-regulation of *PI-KU* by *T. evansi* on the one hand, and *TPS14* and *PR-1* on the other. This could be explained if *T. evansi* secretes different effectors that could affect the plant in different ways, which was shown by Jonckheere et al. (2016).

The beneficial effect of distal *T. evansi* feeding on *T. urticae* oviposition when sharing a leaf has also been observed in other plant species such as tomato where, in contrast to bean, also suppression of defense genes in the distal leaflet areas was observed (Alba et al., 2015). Although this effect appeared to be also sensitive to the timing of infestation (B. C. Schimmel et al., 2017; B. C. J. Schimmel et al., 2017). It is possible that this difference between tomato and bean can be explained by the fact that tomato leaflets are generally smaller in size than bean leaves.

It is possible that the marker genes we used are more suitable for monitoring local responses than for systemic responses. Furthermore, it would be interesting to investigate whether defensive metabolites, such as flavonoids– which are massively induced by *T. urticae*, but not by *T. evansi* (**Table 3**)–are downregulated distally from the *T. evansi* feeding site as a possible explanation for improved *T. urticae* performance on shared leaves (Su et al., 2020). Isovitexin and Genistein 7-O-beta-D-glucoside accumulate to high levels when the plant is treated with phytohormones. *T. urticae* feeding causes an accumulation to similar levels, whereas *T. evansi* feeding does not. The effect of these compound would be interesting to follow up on as potential defensive compounds against mites, or as indirect targets of suppression by *T. evansi*.

### 4.5. Both mite species have a preferred feeding sites and T. evansi displaces T. urticae

Both spider mite species showed preferences for feeding sites on bean leaves. Even though all mites were released on the adaxial leaf surface almost all mites moved to the abaxial side for ovipositing and feeding. On the abaxial side, both species preferred feeding near the petiole and around the midvein (**Fig. 7**). This part of the leaf is the least flat (i.e. veins stick out of the leaf surface) and allows mites to create three-dimensional web structures. This area could also be attractive if nutrients are accumulating more, or defensive compounds accumulate less, in this area but we have not tested for that. We observed that *T. evansi* behaves more gregarious than *T. urticae*, which shows a preference too but is spread out more than *T. evansi*. This behaviour corresponds with earlier studies on the distribution of *T. evansi* and *T. urticae* mites on tomato (Azandémè-Hounmalon et al., 2014).When both mites are present on the same leave at the same time, *T. evansi* does not change its behavior. *T. urticae* mites, however, feed at sites further away from the petiole (**Fig. 7**), indicating that they are either displaced by *T. evansi* or that they avoid their competitors. This behaviour has been shown before on tomato (Godinho et al., 2020). This exclusion by *T. evansi* could be initiated through chemical and/or physical cues. Schimmel et al. (2017) described hyper-suppression of defenses by *T. evansi* in response to the presence of *T. urticae* on the same leave. It is possible that in combination with this changed behavior, *T. evansi* could also be more aggressive and physically attack or push competitors away. This has not been observed, but would be interesting to investigate.

The behavior of *T. evansi* and *T. urticae* appears similar to the behaviour of another spider mite pair observed, *T. neocaledonicus* and *T. ludeni* (Kaimal & Ramani, 2011), albeit in an opposite way. *Tetranychus ludeni*—a defense suppressor (P. Godinho et al., 2016)—was observed to spread over the leaf area at a fast rate whereas *T. neocaledonicus*—susceptible to plant defences (França et al., 2018)—restricted itself to a small area of the leaf and then gradually disperses to other parts of the leaf. *T. evansi* and *T. neocaledonicus* could be clustering as a defense mechanism against predators as it allows for a denser web to be formed faster, increasing protection for the eggs that are laid. At least for *T. evansi* it is known that adult females produce a denser web in response to cues from nearby competitors, which is not the case for *T. urticae* (Sarmento et al., 2011b). Female *T. evansi* mites are also known to prefer using existing web, preferring web of conspecifics, and to share web with other females (Sato et al., 2016). Blaazer et al. (2018) described such buffering traits that allow *T. evansi* to not be outcompeted, despite it suppressing defenses and making the host more attractive to competitors. This study shows that *T. evansi* is able to exclude competitors from its feeding site. The mechanism by which *T. evansi* excludes competitors is not known. Because *T. evansi* and *T. ludeni* are both plant defense supressors, but show opposite gregarious behavior, it is likely this behavior does not relate to dealing with plant defenses. This behavior could have evolved independently of the defense suppression trait in order to deal with competitors or predators.

### 4.6. Conclusion

It seems that in terms of defense manipulation and the effect on competitor oviposition *T. evansi* acts on bean similarly as on tomato. The suppression of gene expression, as made clear by the RNA-seq in this study and the results from Schimmel et al (2018), appears to inhibit the plant response in a transcriptome wide manner, probably by targeting a key regulator for defense. Since this suppression happens in Fabaceae and Solanaceae it is likely to occur in other hosts too. We expect this targeted regulator to be preserved among distinct plant species— considering that *T. evansi* occurs on 35 other families (138 host species in total)—and that the suppression of defenses will therefore be effective on other species as well. We focused out observation on defenses related to or regulated by JA and SA. But considering the transcriptome wide response, the mites might affect the plants on multiple levels, such as manipulation of the metabolism affecting availability of nutrients. We further only briefly investigated mite behaviour in the form of localization. But detailed studies could uncover a more complex interaction between the mite species. We at least show that suppression is likely to occur locally. That can then explain why *T. evansi* has the need to exclude competitors from its feeding site. We therefor recommend further study on defense suppression by *T. evansi* to include other plant families.

## Acknowlegdements

We thank Dr. Patrick Meirmans for his input on analysing the spatial distribution of the mites, and Sonia Jillings for her help with experiments. This research was (partially) funded by NWO Earth and Life Sciences (ALW) project 834.12.003 and 19391 (NWO-VICI) of M.R.K.

## Author contributions

R.C., L.M.S.A., A.S., J.M.A. and M.R.K. conceived and designed the experiments; R.C., L.M.S.A., A.S. and J.M.A. conducted the experiments; data were analysed by R.C., L.M.S.A., B.C.J.S. and M.R.K; R.C. and E.V.P. wrote the manuscript with input from B.C.J.S. and M.R.K. All authors approved the final version of the work.

