## Supplementary material for "Competitor Displacement by an Herbivore that Manipulates Plant Defences"

### Supplemental Figures and Tables

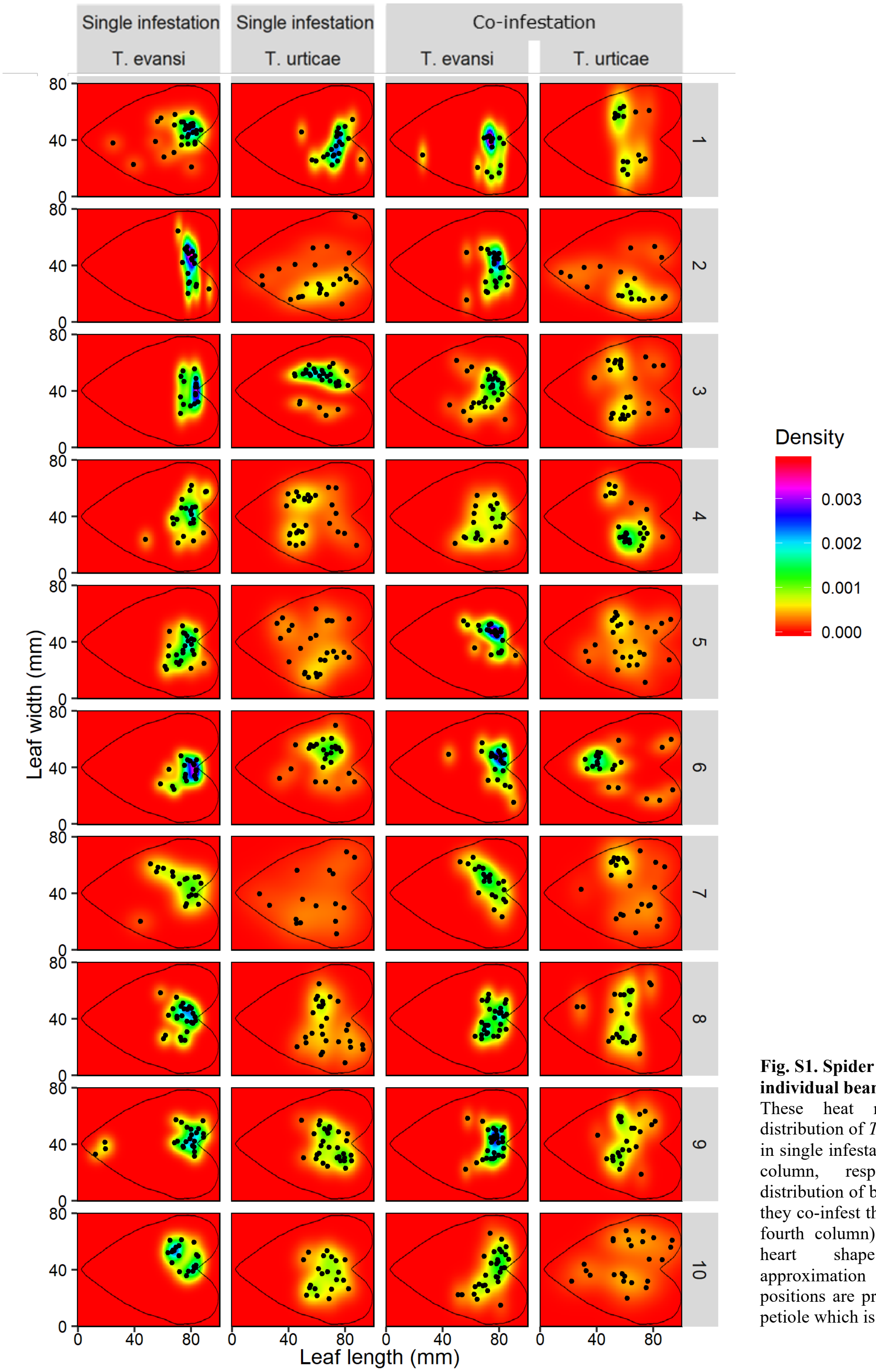


**Fig. S1. Spider mite distribution per individual bean leaf**

These heat maps represent the distribution of *T. evansi* and *T. urticae* in single infestations (first and second column, respectively) and the distribution of both mite species when they co-infest the same leaf (third and fourth column) after 48 hours. The heart shape represents an approximation of a leaf shape. All positions are projected relative to the petiole which is at x: 85; y: 40.

**Table S1.** Bean genes whose expression is **up-regulated by JA but not by SA**. Annotations are based on the refence loci from Arabidopsis or tomato. The 4 most-right columns show the fold change (FC) values of plants treated with salicylic acid (SA), jasmonic acid (JA), *T. urticae* (Tu) or *T. evansi* (Te), relative to control. These genes were **selected based on** two criteria: (1) expression is at least 10 times up-regulated by JA; (2) expression is not significantly upregulated by SA. This table is ranked according to the relative expression level in the jasmonic acid treatment (JA).

| # | Bean locus ID | REFERENCE | ANNOTATION | LOCI | INVOLVED IN | LOCALIZATION | FC expression, relative to control | | | |
|---|---|---|---|---|---|---|---|---|---|---|
| | | | | | | | SA | JA | Tu | Te |
| 1 | Phvul.011G167000.1 | AT5G24090 | Acidic chitinase | | defense response | extracellular region | 0,5 | 496,8 | 2,9 | 3,0 |
| 2 | Phvul.010G135700.1 | AT1G55020 | Lipoxygenase 1 | 4 | defense response | plastid | 1,4 | 383,2 | 12,9 | 3,4 |
| 3 | Phvul.007G276500.1 | Solyc02g087580 | Actin cross-linking | | unknown | cytoplasm | 3,6 | 309,5 | 5,7 | 5,9 |
| 4 | Phvul.008G229400.1 | AT5G59100 | Subtilisin-like serine endopeptidase family protein | 5 | proteolysis | extracellular region | 1,7 | 228,6 | 4,6 | 1,6 |
| 5 | Phvul.003G047100.1 | AT2G15490 | UDP-glycosyltransferase 73B4 | | toxin catabolic process | cytoplasm | 3,0 | 197,8 | 1,2 | 1,1 |
| 6 | Phvul.004G129800.1 | AT1G17860 | Kunitz family trypsin and protease inhibitor protein | 4 | response to stress | extracellular region | 3,2 | 188,0 | 25,1 | 9,0 |
| 7 | Phvul.007G276400.1 | Solyc07g038130 | Beta-galactosidase | | cell wall degradation | extracellular region | 1,6 | 160,4 | 1,7 | 2,6 |
| 8 | Phvul.005G001000.1 | AT4G24010 | Cellulose synthase like G1 | | polysaccharide biosynthetic process | plasma membrane | 1,2 | 159,4 | 0,2 | 0,6 |
| 9 | Phvul.011G143200.1 | AT5G23960 | Terpene synthase 21 | 3 | sesquiterpene biosynthetic process | chloroplast | 0,9 | 131,9 | 1,9 | 3,1 |
| 10 | Phvul.011G000200.1 | AT3G14690 | Cytochrome P450, family 72, subfamily A, polypeptide 15 | | oxygen binding | cytoplasm | 1,1 | 106,5 | 0,3 | 0,6 |
| 11 | Phvul.006G208300.1 | AT2G22590 | UDP-Glycosyltransferase superfamily protein | | unknown | unknown | 0,6 | 89,9 | 0,2 | 0,9 |
| 12 | Phvul.005G032600.1 | AT1G65870 | Disease resistance (dirigent-like) family protein | | defense response | extracellular region | 0,4 | 84,1 | 0,1 | 0,3 |
| 13 | Phvul.007G276600.1 | Solyc06g075020 | ABC transporter G family-like protein | | transport | membrane | 1,1 | 67,9 | 0,9 | 0,9 |
| 14 | Phvul.001G156900.1 | AT1G06800 | alpha/beta-Hydrolases superfamily protein | | lipid metabolic process | chloroplast | 3,5 | 63,1 | 7,9 | 2,6 |
| 15 | Phvul.010G165100.1 | AT1G78950 | Terpenoid cyclases family protein | | sterol biosynthetic process | nucleus | 0,7 | 48,6 | 0,1 | 0,4 |
| 16 | Phvul.009G116300.1 | AT3G12500 | Basic chitinase | | defense response | vacuole | 1,6 | 45,9 | 3,1 | 1,4 |
| 17 | Phvul.006G034000.1 | AT1G78570 | Rhamnose biosynthesis 1 | | flavonol biosynthetic process | cytoplasm | 0,9 | 45,1 | 0,5 | 0,5 |
| 18 | Phvul.011G107200.1 | AT3G25830 | Terpene synthase-like sequence-1,8-cineole | | monoterpenoid biosynthetic process | chloroplast | 0,8 | 39,0 | 1,3 | 4,2 |
| 19 | Phvul.010G158400.1 | AT4G37850 | Basic helix-loop-helix (bHLH) DNA-binding protein | | transcription | nucleus | 3,0 | 37,9 | 1,8 | 1,1 |
| 20 | Phvul.002G216900.1 | AT1G58440 | FAD/NAD(P)-binding oxidoreductase family protein | | response to stress | extracellular region | 2,9 | 37,5 | 0,2 | 0,5 |
| 21 | Phvul.007G277100.1 | Solyc03g065340 | Alpha-1,4 glucan phosphorylase | | carbohydrate metabolism. | cytoplasm | 0,9 | 30,9 | 1,1 | 0,9 |
| 22 | Phvul.005G111500.1 | AT4G16730 | Terpene synthase 02 | | monoterpenoid biosynthetic process | chloroplast | 2,1 | 25,0 | 1,6 | 1,0 |
| 23 | Phvul.003G170600.1 | AT5G34850 | Purple acid phosphatase 26 | | phosphate ion homeostasis | vacuole | 0,8 | 21,3 | 1,1 | 1,1 |
| 24 | Phvul.011G107000.1 | AT4G03210 | Xyloglucan endotransglucosylase/hydrolase 9 | | regulation of meristem growth | extracellular region | 1,3 | 21,2 | 0,3 | 0,6 |
| 25 | Phvul.010G013000.1 | AT4G37330 | Cytochrome P450, family 81, subfamily D, polypeptide 4 | | oxygen binding | endoplasmic reticulum | 2,4 | 20,7 | 15,5 | 1,6 |
| 26 | Phvul.003G287500.1 | AT4G37970 | Cinnamyl alcohol dehydrogenase 6 | | lignin biosynthetic process | cytoplasm | 0,9 | 19,9 | 1,5 | 1,1 |
| 27 | Phvul.003G076800.1 | AT1G61820 | Beta glucosidase 46 | | lignin biosynthetic process | extracellular region | 1,4 | 18,3 | 0,5 | 0,2 |
| 28 | Phvul.003G052000.1 | AT1G23760 | BURP domain-containing protein | | xylan biosynthetic process | extracellular region | 0,7 | 14,6 | 0,1 | 0,5 |
| 29 | Phvul.011G113000.1 | AT4G03400 | Auxin-responsive GH3 family protein | | response to stress | chloroplast | 2,2 | 14,0 | 0,6 | 1,0 |
| 30 | Phvul.010G012900.1 | AT4G37370 | Cytochrome P450, family 81, subfamily D, polypeptide 8 | 2 | response to stress | endoplasmic reticulum | 2,0 | 13,9 | 9,5 | 1,4 |
| 31 | Phvul.011G197000.1 | AT5G16080 | Carboxyesterase 17 | | unknown | unknown | 1,6 | 11,0 | 9,2 | 7,5 |
| 32 | Phvul.002G016400.1 | AT2G15480 | UDP-glucosyl transferase 73B5 | 2 | toxin catabolic process | unknown | 3,1 | 10,6 | 0,9 | 1,4 |
| 33 | Phvul.006G195600.1 | AT1G61680 | Terpene synthase 14 | | monoterpene biosynthetic process | chloroplast | 1,0 | 9,8 | 5,2 | 1,8 |
| 34 | Phvul.008G250700.1 | AT4G35160 | O-methyltransferase family protein | | phenylpropanoid metabolic process | cytoplasm | 2,9 | 8,9 | 31,0 | 8,1 |

Colored cells are significantly different compared to the control treatment, adjusted p-value<0.05

**Table S2.** Bean genes whose expression is **up-regulated by SA but not by JA**. Annotations are based on the refence loci from Arabidopsis or tomato. The 4 most-right columns show the fold change (FC) values of plants treated with salicylic acid (SA), jasmonic acid (JA), *T. urticae* (Tu) or *T. evansi* (Te), relative to control. These genes were **selected based on** two criteria: (1) expression is at least 4 times up-regulated by SA and (2) is not significantly upregulated by JA. This table is ranked according to the relative expression level in the salicylic acid treatment (SA).

| | | | | | | FC expression, relative to control | | | |
|---|---|---|---|---|---|---|---|---|---|
| **#** | **Bean locus ID** | **REFERENCE** | **ANNOTATION** | **INVOLVED IN** | **LOCALIZATION** | **SA** | **JA** | **Tu** | **Te** |
| 1 | Phvul.001G128500.1 | AT3G57270 | Beta-1,3-glucanase 1 | unknown | extracellular region | 42,0 | 0,7 | 1124,1 | 10,6 |
| 2 | Phvul.005G163500.1 | AT1G51800 | Leucine-rich repeat protein kinase family protein | defense response | plasma membrane | 24,8 | 0,8 | 200,0 | 23,8 |
| 3 | Phvul.002G204500.1 | AT1G25270 | Nodulin MtN21 /EamA-like transporter family protein | unknown | extracellular region | 23,6 | 1,2 | 56,6 | 1,1 |
| 4 | Phvul.005G164000.1 | AT1G51800 | Leucine-rich repeat protein kinase family protein | defense response | plasma membrane | 21,8 | 1,3 | 188,7 | 13,4 |
| 5 | Phvul.008G232700.1 | AT3G55470 | Calcium-dependent lipid-binding (CaLB domain) family protein | positive regulation of flavonoid biosynthesis | unknown | 18,9 | 1,6 | 248,0 | 6,8 |
| 6 | Phvul.004G105100.1 | AT2G34930 | Disease resistance family protein / LRR family protein | defense response | cell wall | 17,3 | 1,0 | 104,5 | 9,1 |
| 7 | Phvul.001G039900.1 | AT5G26170 | WRKY 50 | jasmonic acid mediated signaling pathway | nucleus | 14,0 | 0,6 | 304,0 | 11,0 |
| 8 | Phvul.004G142500.1 | AT2G25470 | Receptor like protein 21 | signal transduction | chloroplast | 13,0 | 0,9 | 110,9 | 11,3 |
| 9 | Phvul.011G177000.1 | AT4G16260 | Glycosyl hydrolase superfamily protein | defense response | membrane | 12,2 | 1,9 | 329,9 | 4,8 |
| 10 | Phvul.001G040300.1 | AT5G10530 | L-type lectin receptor kinase IX.1 | defense response | extracellular region | 11,0 | 1,0 | 235,3 | 8,8 |
| 11 | Phvul.010G021700.1 | AT4G27860 | Vacuolar iron transporter (VIT) family protein | iron ion transmembrane transporter | endoplasmic reticulum | 8,8 | 1,0 | 450,1 | 59,0 |
| 12 | Phvul.005G009500.1 | AT5G61520 | Major facilitator superfamily protein | carbohydrate transmembrane transporter | membrane | 8,2 | 1,5 | 13,8 | 2,8 |
| 13 | Phvul.002G286500.1 | AT4G11650 | Osmotin 34 | defense response | extracellular region | 7,8 | 0,8 | 1175,2 | 10,7 |
| 14 | Phvul.005G087100.1 | AT4G27220 | NB-ARC domain-containing disease resistance protein | defense response | nucleus | 6,9 | 1,0 | 81,3 | 7,9 |
| 15 | Phvul.005G027000.1 | Solyc01g010393 | Beta-glucosidase | response to stress | extracellular region | 6,6 | 1,8 | 23,5 | 5,2 |
| 16 | Phvul.005G173700.1 | AT1G15520 | Pleiotropic drug resistance 12 | defense response | plasma membrane | 6,5 | 1,9 | 1,8 | 1,0 |
| 17 | Phvul.008G044400.1 | AT4G08850 | Leucine-rich repeat receptor-like protein kinase family protein | response to stress | membrane | 6,3 | 1,9 | 48,4 | 3,7 |
| 18 | Phvul.008G045200.1 | AT4G08850 | Leucine-rich repeat receptor-like protein kinase family protein | response to stress | membrane | 6,0 | 1,8 | 49,9 | 2,8 |
| 19 | Phvul.008G007900.1 | AT1G66880 | Protein kinase superfamily protein | defense response | plasma membrane | 5,9 | 1,9 | 45,7 | 2,1 |
| 20 | Phvul.006G007500.1 | AT5G47650 | Nudix hydrolase homolog 2 | response to oxidative stress | cytoplasm | 5,8 | 1,9 | 26,4 | 3,1 |
| 21 | Phvul.001G183800.1 | AT2G36670 | Eukaryotic aspartyl protease family protein | proteolysis | extracellular region | 5,7 | 1,8 | 11,6 | 2,6 |
| 22 | Phvul.007G264500.1 | AT5G06600 | Ubiquitin-specific protease 12 | glucose catabolic process | cytoplasm | 5,5 | 1,5 | 23,0 | 2,3 |
| 23 | Phvul.003G205900.1 | AT4G24730 | Calcineurin-like metallo-phosphoesterase superfamily protein | unknown | plasma membrane | 5,1 | 1,6 | 13,0 | 3,4 |
| 24 | Phvul.005G030700.1 | AT3G27890 | NADPH:quinone oxidoreductase | defense response | chloroplast | 4,8 | 1,9 | 3,1 | 1,6 |
| 25 | Phvul.010G016700.1 | AT2G37770 | NAD(P)-linked oxidoreductase superfamily protein | response to stress | cytoplasm | 4,6 | 1,6 | 4,1 | 1,3 |
| 26 | Phvul.005G057500.1 | AT2G32190 | Cysteine-rich transmembrane module 4 | response to chitin | plasma membrane | 4,3 | 1,9 | 3,1 | 1,2 |
| 27 | Phvul.011G179500.1 | AT1G79380 | Ca(2)-dependent phospholipid-binding protein (Copine) | defense response | plasma membrane | 4,2 | 1,0 | 22,9 | 1,2 |
| 28 | Phvul.008G273600.1 | AT1G45616 | Receptor like protein 6 | defense response | membrane | 4,1 | 1,7 | 14,9 | 2,9 |
| 29 | Phvul.002G171400.1 | AT5G17680 | Disease resistance protein (TIR-NBS-LRR class) | defense response | unknown | 4,1 | 1,5 | 8,7 | 2,2 |
| 30 | Phvul.003G268600.1 | AT3G12500 | Basic chitinase | defense response | vacuole | 4,1 | 1,4 | 7,0 | 1,6 |

Colored cells are significantly different compared to the control treatment, adjusted p-value<0.05

**Table S3.** Bean genes whose expression is **down-regulated by JA**. Annotations are based on the refence loci from Arabidopsis or tomato. The 4 most-right columns show the fold change (FC) values of plants treated with salicylic acid (SA), jasmonic acid (JA), *T. urticae* (Tu) or *T. evansi* (Te), relative to control. These genes were **selected based on** the following criteria: expression is significantly down-regulated in the JA sample, but not in the SA sample. This table is ranked according to the relative expression level in the salicylic acid treatment (SA).

| | | | | | | FC expression, relative to control | | | |
|---|---|---|---|---|---|---|---|---|---|
| **#** | **Bean locus ID** | **REFERENCE** | **ANNOTATION** | **INVOLVED IN** | **LOCALIZATION** | **SA** | **JA** | **Tu** | **Te** |
| 1 | Phvul.006G004900.1 | AT1G65610 | Six-hairpin glycosidases superfamily protein | unknown | plasma membrane | 4,9 | DOWN | 235,4 | 10,3 |
| 2 | Phvul.005G162400.1 | AT2G19210 | Leucine-rich repeat transmembrane protein kinase protein | unknown | plasma membrane | 4,5 | DOWN | 41,3 | 2,1 |
| 3 | Phvul.005G163100.1 | AT1G51800 | Leucine-rich repeat protein kinase family protein | defense response | plasma membrane | 4,4 | DOWN | 379,4 | 3,3 |
| 4 | Phvul.005G077600.1 | AT3G14840 | Leucine-rich repeat transmembrane protein kinase | protein phosphorylation | plasma membrane | 2,8 | DOWN | 31,9 | 1,3 |
| 5 | Phvul.005G164100.1 | AT4G29990 | Leucine-rich repeat transmembrane protein kinase protein | protein phosphorylation | plasma membrane | 2,2 | DOWN | 207,8 | 2,7 |
| 6 | Phvul.008G106500.1 | AT4G08850 | Leucine-rich repeat receptor-like protein kinase | response to stress | membrane | 2,0 | DOWN | 10,4 | 1,6 |
| 7 | Phvul.004G019000.1 | AT3G01420 | Peroxidase superfamily protein | cell death | extracellular region | 1,9 | DOWN | 8,8 | DOWN |
| 8 | Phvul.005G162800.1 | Solyc01g108900 | Pollen leucine rich repeat domain-extensin | pollen tube growth | extracellular region | 1,8 | DOWN | 67,8 | 2,5 |
| 9 | Phvul.001G087500.1 | AT5G14020 | Endosomal targeting BRO1-like domain-containing protein | unknown | vacuole | 1,8 | DOWN | 1,0 | DOWN |
| 10 | Phvul.010G045100.1 | Solyc09g083220 | bHLH transcription factor 60 | response to stress | extracellular region | 1,7 | DOWN | 42,7 | 1,3 |
| 11 | Phvul.007G186900.1 | AT4G14746 | Neurogenic locus notch-like protein | guard cell | membrane | 1,6 | DOWN | 289,9 | 1,0 |
| 12 | Phvul.L006200.1 | AT2G34930 | Disease resistance family protein / LRR family protein | defense response | cell wall | 1,6 | DOWN | 23,1 | 2,0 |
| 13 | Phvul.006G173200.1 | Solyc07g006420 | Transmembrane protein | unknown | membrane | 1,5 | DOWN | 43,6 | 1,6 |
| 14 | Phvul.002G199700.1 | AT5G44400 | FAD-binding Berberine family protein | unknown | cell wall | 1,4 | DOWN | 10,8 | DOWN |
| 15 | Phvul.006G003400.1 | AT2G34930 | Disease resistance family protein / LRR family protein | defense response | cell wall | 1,3 | DOWN | 39,0 | 1,2 |
| 16 | Phvul.001G028100.1 | AT1G59590 | ZCF37 | unknown | mitochondrion | 1,3 | 0,2 | 3,8 | 0,5 |
| 17 | Phvul.010G012700.1 | AT2G45550 | Cytochrome P450, family 76, subfamily C, polypeptide 4 | oxygen binding | chloroplast | 1,1 | DOWN | 418,8 | 2,5 |
| 18 | Phvul.002G017500.1 | AT4G34050 | S-adenosyl-L-methionine-dependent methyltransferase | phenylpropanoid metabolism | cytoplasm | 1,0 | DOWN | 26,8 | 2,4 |
| 19 | Phvul.002G128500.1 | AT5G45390 | CLP protease P4 | photosystem II assembly | chloroplast | 1,0 | 0,7 | 0,7 | 0,8 |
| 20 | Phvul.010G081700.1 | AT2G45270 | Glycoprotease 1 | proteolysis | mitochondrion | 0,9 | 0,8 | 0,7 | 0,7 |

Colored cells are significantly different compared to the control treatment, adjusted p-value<0.05

**Table S4.** Bean genes whose expression is **down-regulated by SA**. Annotations are based on the refence loci from Arabidopsis or tomato. The 4 most-right columns show the fold change (FC) values of plants treated with salicylic acid (SA), jasmonic acid (JA), *T. urticae* (Tu) or *T. evansi* (Te), relative to control. These genes were **selected based on** the following criteria: expression is significantly down-regulated in the SA sample, but not in the JA sample. This table is ranked according to the relative expression level in the jasmonic acid treatment (JA).

| | | | | | | FC expression, relative to control | | | |
|---|---|---|---|---|---|---|---|---|---|
| **#** | **Bean locus ID** | **REFERENCE** | **ANNOTATION** | **INVOLVED IN** | **LOCALIZATION** | **SA** | **JA** | **Tu** | **Te** |
| 1 | Phvul.010G165000.1 | AT1G16760 | Protein kinase | plant-type cell wall modification | nucleus | DOWN | 17,6 | DOWN | DOWN |
| 2 | Phvul.011G044900.1 | AT4G39250 | RAD-like 1 | transcription | nucleus | DOWN | 5,4 | DOWN | 1,4 |
| 3 | Phvul.011G142600.1 | AT5G23960 | Terpene synthase 21 | sesquiterpene biosynthesis | chloroplast | DOWN | 3,3 | 1,3 | DOWN |
| 4 | Phvul.010G101400.1 | AT5G33406 | hAT dimerisation domain-containing protein | transposase-like protein | unknown | DOWN | 1,2 | 4,8 | 0,8 |
| 5 | Phvul.011G105600.1 | AT2G46410 | Homeodomain-like superfamily protein | epidermal cell differentiation | nucleus | DOWN | 1,0 | 27,5 | DOWN |
| 6 | Phvul.008G020900.1 | AT1G12210 | RPS5-like 1 | defense response | unknown | DOWN | 1,0 | 8,4 | 1,0 |
| 7 | Phvul.001G040800.1 | AT5G10530 | L-type lectin receptor kinase IX.1 | defense response | extracellular region | DOWN | 0,8 | 29,9 | 0,8 |
| 8 | Phvul.011G056600.1 | AT3G57880 | Calcium-dependent lipid-binding family protein | unknown | endoplasmic reticulum | DOWN | 0,7 | DOWN | 0,7 |

Colored cells are significantly different compared to the control treatment, adjusted p-value<0.05

**Table S5.** Bean genes whose expression is **up-regulated by both JA and SA**. Annotations are based on the refence loci from Arabidopsis or tomato. The 5 most-right columns show the fold change (FC) values of plants treated with salicylic acid (SA), jasmonic acid (JA), *T. urticae* (Tu) or *T. evansi* (Te), relative to control or *T. urticae* relative to *T. evansi* (Tu/Te). These genes were **selected based on** one criterium, i.e. expression is significantly up-regulated in the JA sample and in the SA sample. This table is ranked according to the relative difference in expression between the *T. urticae* and *T. evansi* treatment (Tu/Te).

| # | Bean locus ID | REFERENCE | ANNOTATION | LOCI | INVOLVED IN | LOCALIZATION | FC expression, relative to control | | | | |
|---|---|---|---|---|---|---|---|---|---|---|---|
| | | | | | | | SA | JA | Tu | Te | Tu/Te |
| 1 | Phvul.002G285800.1 | AT5G13080 | WRKY DNA-binding protein 75 | | response to stress | nucleus | UP | UP | UP | NA | UP |
| 2 | Phvul.004G130000.1 | AT1G17860 | Kunitz family trypsin inhibitor | | response to stress | extracellular region | UP | UP | UP | NA | UP |
| 3 | Phvul.006G079600.1 | AT1G78370 | Glutathione S-transferase TAU 20 | | toxin catabolic process | apoplast | UP | UP | UP | NA | UP |
| 4 | Phvul.007G211500.1 | Solyc01g100940 | Starch synthase 2 | | starch biosynthesis | chloroplast | UP | UP | UP | NA | UP |
| 5 | Phvul.011G203600.1 | Solyc07g054720 | Type II proteinase inhibitor family protein | | proteolysis | extracellular region | UP | UP | UP | NA | UP |
| 6 | Phvul.007G049700.1 | AT4G21410 | Cysteine-rich RECEPTOR-like protein kinase 29 | | response to abscisic acid | plasma membrane | UP | UP | UP | UP | 510,1 |
| 7 | Phvul.002G038700.1 | AT5G13930 | Chalcone and stilbene synthase family protein | 3 | flavonoid biosynthetic process | vacuole | UP | UP | UP | UP | 442,0 |
| 8 | Phvul.010G120200.1 | AT5G54160 | O-methyltransferase 1 | 2 | phenylpropanoid metabolism | cytoplasm | UP | UP | UP | UP | 313,9 |
| 9 | Phvul.003G192000.1 | AT5G15130 | WRKY DNA-binding protein 72 | | defense response | nucleus | UP | UP | UP | UP | 78,0 |
| 10 | Phvul.005G054100.1 | AT1G78380 | Glutathione S-transferase TAU 19 | 2 | toxin catabolic process | chloroplast | UP | UP | UP | UP | 69,8 |
| 11 | Phvul.005G080300.1 | AT3G56400 | WRKY DNA-binding protein 70 | 2 | defense response | nucleus | UP | UP | UP | UP | 57,8 |
| 12 | Phvul.002G317000.1 | AT1G25340 | MYB domain protein 116 | | phenylpropanoid metabolism | nucleus | 175,9 | 129,7 | 63,8 | 2,6 | 24,5 |
| 13 | Phvul.006G056600.1 | Solyc01g086810 | NBS-LRR disease resistance protein | | defense response | extracellular region | UP | UP | UP | UP | 18,7 |
| 14 | Phvul.001G019200.1 | AT4G37260 | MYB domain protein 73 | | glucosinolate metabolic process | nucleus | UP | UP | UP | UP | 17,2 |
| 15 | Phvul.006G033300.1 | AT5G60900 | Receptor-like protein kinase 1 | 2 | defense response | plasma membrane | UP | UP | UP | UP | 15,0 |
| 16 | Phvul.005G075500.1 | AT1G17010 | 2-oxoglutarate (2OG) and Fe(II)-dependent oxygenase | | flavonoid biosynthetic process | cytoplasm | UP | UP | UP | UP | 14,9 |
| 17 | Phvul.008G109200.1 | AT1G58190 | Receptor like protein 9 | | signal transduction | extracellular region | UP | UP | UP | UP | 9,6 |
| 18 | Phvul.011G189300.1 | AT1G76680 | 12-oxophytodienoate reductase 1 | | response to wounding | cytoplasm | UP | UP | UP | UP | 9,4 |
| 19 | Phvul.011G056100.1 | AT5G44640 | Beta glucosidase 13 | | glucosinolate catabolic process | Golgi apparatus | UP | UP | UP | UP | 8,8 |
| 20 | Phvul.002G214900.1 | AT5G10530 | L-type lectin receptor kinase IX.1 | | defense response | extracellular region | UP | UP | UP | UP | 8,0 |
| 21 | Phvul.007G125500.1 | AT2G38870 | Serine protease inhibitor type I | | defense response | extracellular region | UP | UP | UP | UP | 6,4 |
| 22 | Phvul.002G018100.1 | AT4G36950 | MAPKKK21 | | signaling | plasma membrane | UP | UP | UP | UP | 4,2 |
| 23 | Phvul.011G109600.1 | AT3G23250 | MYB domain protein 15 | | phenylpropanoid metabolism | nucleus | UP | UP | UP | UP | 3,9 |
| 24 | Phvul.008G058400.1 | AT3G61510 | ACC synthase 1 | | ethylene biosynthetic process | cytoplasm | UP | UP | UP | UP | 3,6 |
| 25 | Phvul.011G189200.1 | AT1G76690 | 12-oxophytodienoate reductase 2 | | response to wounding | cytoplasm | UP | UP | UP | UP | 2,7 |
| 26 | Phvul.002G014400.1 | AT4G37390 | Auxin-responsive GH3 family protein | | auxin homeostasis | cytoplasm | 239,1 | 218,6 | 1,9 | 1,0 | 2,0 |
| 27 | Phvul.002G144800.1 | AT4G37990 | Elicitor-activated gene 3-2 | | plant-type HR response | cytoplasm | UP | UP | UP | UP | 1,9 |
| 28 | Phvul.007G077800.1 | AT5G24090 | Acidic chitinase | | defense response | extracellular region | UP | UP | UP | UP | 1,6 |
| 29 | Phvul.006G069500.1 | AT2G30020 | Protein phosphatase 2C family protein | | defense response | plastid | UP | UP | NA | UP | DOWN |

Colored cells are significantly different compared to the control treatment, adjusted p-value<0.05

**Table S6.** Bean genes whose expression is **down-regulated by both JA and SA**. Annotations are based on the refence loci from Arabidopsis or tomato. The 5 most-right columns show the fold change (FC) values of plants treated with salicylic acid (SA), jasmonic acid (JA), *T. urticae* (Tu) or *T. evansi* (Te), relative to control or *T. urticae* relative to *T. evansi* (Tu/Te). These genes were **selected based on** the criterium that expression is significantly down-regulated in both the JA and the SA sample. This table is ranked according to the relative expression in the *T. urticae* treatment (Tu).

| # | Bean locus ID | REFERENCE | ANNOTATION | LOCI | INVOLVED IN | LOCALIZATION | FC expression, relative to control | | | | |
|---|---|---|---|---|---|---|---|---|---|---|---|
| | | | | | | | SA | JA | Tu | Te | Tu/Te |
| 1 | Phvul.009G103800.1 | AT4G31320 | SAUR-like auxin-responsive protein family | | response to auxin | nucleus | DOWN | DOWN | 26,6 | DOWN | UP |
| 2 | Phvul.007G064500.1 | AT1G52540 | Protein kinase superfamily protein | | protein phosphorylation | plasma membrane | DOWN | DOWN | 16,0 | 1,0 | 16,0 |
| 3 | Phvul.007G228100.1 | AT2G18950 | Homogentisate phytyltransferase 1 | | defense response | plasma membrane | DOWN | DOWN | 12,5 | 1,0 | 12,5 |
| 4 | Phvul.002G008000.1 | Solyc04g008700 | Transcription elongation regulator 1 | | transcription | nucleus | DOWN | DOWN | 7,6 | DOWN | UP |
| 5 | Phvul.009G121200.1 | AT1G77740 | Phosphatidylinositol-4-phosphate 5-kinase 2 | | growth | plasma membrane | 0,5 | 0,5 | 0,4 | 0,7 | 0,5 |

Colored cells are significantly different compared to the control treatment, adjusted p-value<0.05

**Table S7.** Bean genes whose expression is **up-regulated by Tu but not by Te**. Annotations are based on the refence loci from Arabidopsis or tomato. The 4 most-right columns show the fold change (FC) values of plants treated with salicylic acid (SA), jasmonic acid (JA), *T. urticae* (Tu) or *T. evansi* (Te), relative to control. These genes were **selected based on** two criteria: (1) expression is significantly up-regulated in the Tu sample; (2) expression is not significantly up-regulated in by Te. This table is ranked according to the relative expression level in the *T. urticae* treatment (Tu).

| | | | | | | | FC expression, relative to control | | | |
|---|---|---|---|---|---|---|---|---|---|---|
| # | Bean locus ID | REFERENCE | ANNOTATION | LOCI | INVOLVED IN | LOCALIZATION | SA | JA | Tu | Te |
| 1 | Phvul.004G021400.1 | AT4G31940 | Cytochrome P450, family 82, subfamily C, polypeptide 4 | 2 | response to stress | membrane | 3,6 | 1,3 | 644,6 | 2,7 |
| 2 | Phvul.002G033000.1 | AT4G39230 | NmrA-like negative transcriptional regulator | 2 | response to cadmium ion | cytoplasm | 11,0 | 2,8 | 598,4 | 1,5 |
| 3 | Phvul.007G091000.1 | AT5G54160 | O-methyltransferase 1 | | phenylpropanoid metabolism | cytoplasm | 3,8 | 3,9 | 494,0 | 2,4 |
| 4 | Phvul.002G144600.1 | AT4G37990 | Elicitor-activated gene 3-2 | | plant-type HR response | cytoplasm | 122,2 | 2,0 | 282,6 | 1,0 |
| 5 | Phvul.006G188500.1 | Solyc02g078650 | Polyphenol oxidase | | defense response | chloroplast | 348,4 | 152,6 | 247,9 | 2,0 |
| 6 | Phvul.010G012800.1 | AT3G62150 | P-glycoprotein 21 | | auxin efflux | membrane | 1,6 | 0,6 | 222,4 | 1,9 |
| 7 | Phvul.004G018900.1 | AT3G01420 | Peroxidase superfamily protein | | cell death | extracellular region | 11,6 | 6,1 | 219,5 | 2,7 |
| 8 | Phvul.005G164100.1 | AT4G29990 | Leucine-rich repeat transmembrane protein kinase | | protein phosphorylation | plasma membrane | 2,2 | DOWN | 207,8 | 2,7 |
| 9 | Phvul.010G115500.1 | AT5G49330 | MYB domain protein 111 | | flavonol biosynthetic process | nucleus | 6,7 | 3,3 | 207,1 | 1,2 |
| 10 | Phvul.006G074600.1 | AT2G38470 | WRKY DNA-binding protein 33 | | defense response | nucleus | 50,1 | 8,8 | 198,1 | 1,3 |
| 11 | Phvul.011G214400.1 | AT5G60900 | Receptor-like protein kinase 1 | 3 | defense response | plasma membrane | 3,5 | 2,0 | 195,1 | 1,4 |
| 12 | Phvul.008G228700.1 | AT3G46850 | Subtilase family protein | 2 | proteolysis | extracellular region | 1,3 | 1,8 | 184,4 | 2,6 |
| 13 | Phvul.009G244200.1 | AT4G37340 | Cytochrome P450, family 81, subfamily D, polypeptide 3 | | oxygen binding | unknown | 4,2 | 1,2 | 176,1 | 1,0 |
| 14 | Phvul.002G149300.1 | AT5G65380 | MATE efflux family protein | | fruit ripening | plasma membrane | 33,4 | 17,7 | 165,7 | 3,0 |
| 15 | Phvul.007G228000.1 | AT2G18950 | Homogentisate phytyltransferase 1 | | defense response | plasma membrane | 5,0 | 2,0 | 155,5 | 1,7 |
| 16 | Phvul.005G054200.1 | AT1G78380 | glutathione S-transferase TAU 19 | | toxin catabolic process | chloroplast | 17,5 | 11,2 | 142,6 | 1,5 |
| 17 | Phvul.002G199800.1 | AT2G34790 | FAD-binding Berberine family protein | | cell wall organization | cell wall | 4,3 | 2,1 | 139,6 | 1,6 |
| 18 | Phvul.007G242100.1 | AT5G59100 | Subtilisin-like serine endopeptidase family protein | | proteolysis | extracellular region | 3,4 | 2,1 | 138,1 | 2,1 |
| 19 | Phvul.001G131000.1 | AT2G41690 | Heat shock transcription factor B3 | | toxin catabolic process | nucleus | 17,9 | 12,3 | 133,3 | 1,6 |
| 20 | Phvul.009G046900.1 | AT3G47570 | Leucine-rich repeat protein kinase family protein | 2 | protein phosphorylation | plasma membrane | 2,4 | 0,6 | 119,4 | 0,6 |
| 21 | Phvul.001G145600.1 | AT5G42500 | Disease resistance dirigent-like protein | 2 | defense response | extracellular region | 39,5 | 28,0 | 104,6 | 2,4 |
| 22 | Phvul.002G025000.1 | AT5G25120 | Cytochrome p450, family 71, subfamily B, polypeptide 11 | | oxygen binding | plasma membrane | 6,8 | 2,2 | 102,0 | 0,9 |
| 23 | Phvul.004G071700.1 | AT2G39200 | Seven transmembrane MLO family protein | | toxin catabolic process | plasma membrane | 32,3 | 12,0 | 101,4 | 2,3 |
| 24 | Phvul.011G150200.1 | AT4G27290 | S-locus lectin protein kinase family protein | | root hair cell differentiation | plasma membrane | 2,7 | 3,9 | 92,5 | 2,6 |
| 25 | Phvul.002G265400.1 | AT5G64810 | WRKY DNA-binding protein 51 | | jasmonic acid mediated signaling | nucleus | 4,8 | 0,8 | 89,2 | 2,3 |
| 26 | Phvul.011G166500.1 | AT1G17860 | Kunitz family trypsin and protease inhibitor protein | 2 | response to stress | extracellular region | 1,9 | 2,1 | 88,3 | 1,1 |
| 27 | Phvul.003G051800.1 | AT5G06900 | Cytochrome P450, family 93, subfamily D, polypeptide 1 | | oxygen binding | unknown | 139,5 | 77,3 | 87,5 | 0,8 |
| 28 | Phvul.006G124600.1 | AT1G30700 | FAD-binding Berberine family protein | 2 | toxin catabolic process | chloroplast | 11,7 | 6,3 | 82,9 | 1,2 |
| 29 | Phvul.001G140100.1 | AT5G06839 | bZIP transcription factor family protein | | response to xenobiotic stimulus | nucleus | 13,3 | 7,0 | 76,8 | 1,3 |
| 30 | Phvul.004G125200.1 | AT4G05200 | Cysteine-rich RLK (RECEPTOR-like protein kinase) 25 | | defense response | extracellular region | 4,2 | 0,7 | 74,9 | 1,3 |
| 31 | Phvul.006G079700.1 | AT2G30490 | cinnamate-4-hydroxylase | | phenylpropanoid metabolism | vacuolar membrane | 13,3 | 4,0 | 68,1 | 1,9 |
| 32 | Phvul.002G317000.1 | AT1G25340 | MYB domain protein 116 | | regulation of transcription | nucleus | 175,9 | 129,7 | 63,8 | 2,6 |
| 33 | Phvul.006G056500.1 | AT3G07040 | NB-ARC domain-containing disease resistance protein | | plant-type HR response | plasma membrane | 3,5 | 1,5 | 58,1 | 3,0 |

Colored cells are significantly different compared to the control treatment, adjusted p-value<0.05

**Table S8.** Bean genes whose expression is **down-regulated by Tu but not by Te**. Annotations are based on the refence loci from Arabidopsis or tomato. The 4 most-right columns show the fold change (FC) values of plants treated with salicylic acid (SA), jasmonic acid (JA), *T. urticae* (Tu) or *T. evansi* (Te), relative to control. These genes were **selected based on** two criteria: (1) expression is significantly down-regulated in the Tu sample; (2) expression is not significantly down-regulated in by Te. This table is ranked according to the relative expression level in the *T. urticae* treatment (Tu).

| | | | | | | | FC expression, relative to control | | | |
|---|---|---|---|---|---|---|---|---|---|---|
| **#** | **Bean locus ID** | **REFERENCE** | **ANNOTATION** | **LOCI** | **INVOLVED IN** | **LOCALIZATION** | **SA** | **JA** | **Tu** | **Te** |
| 1 | Phvul.003G266300.1 | AT5G66090 | Cell wall integrity/stress response component | | unknown | chloroplast | 0,9 | 0,9 | 0,7 | 0,9 |
| 2 | Phvul.011G004500.1 | AT5G51545 | Low psii accumulation2 | | photosystem II assembly | chloroplast | 0,8 | 0,8 | 0,7 | 0,9 |
| 3 | Phvul.009G186200.1 | AT1G18730 | NDH dependent flow 6 | | photosynthesis | chloroplast | 0,5 | 0,5 | 0,7 | 1,0 |
| 4 | Phvul.009G033600.1 | AT4G09040 | RNA-binding (RRM/RBD/RNP motifs) family protein | | mRNA splicing | chloroplast | 0,8 | 0,8 | 0,7 | 0,9 |
| 5 | Phvul.003G166300.1 | AT4G02530 | Chloroplast thylakoid lumen protein | | response to stress | chloroplast | 0,7 | 0,7 | 0,7 | 0,9 |
| 6 | Phvul.001G055300.1 | AT4G32590 | 2Fe-2S ferredoxin-like superfamily protein | | abscisic acid biosynthetic process | unknown | 0,8 | 0,9 | 0,7 | 0,9 |
| 7 | Phvul.003G229700.1 | AT1G32520 | Oxidation resistance 4 | | unknown | chloroplast | 0,6 | 0,8 | 0,6 | 0,9 |
| 8 | Phvul.004G069700.1 | AT3G26710 | Cofactor assembly of complex C | | photosystem II assembly | chloroplast | 0,6 | 0,6 | 0,6 | 0,9 |
| 9 | Phvul.011G051300.1 | Solyc07g015980 | Purple acid phosphatase | 2 | phosphate ion homeostasis | vacuole | 0,7 | 0,7 | 0,6 | 0,9 |
| 10 | Phvul.002G090400.1 | ATCG00905 | Ribosomal protein S12C | | translation | chloroplast | 1,0 | 0,8 | 0,6 | 1,0 |
| 11 | Phvul.006G209500.1 | AT3G26300 | Cytochrome P450, family 71, subfamily B, polypeptide 34 | | oxygen binding | chloroplast | 0,7 | 0,6 | 0,5 | 0,9 |
| 12 | Phvul.003G022100.1 | AT2G01590 | Chlororespiratory reduction 3 | | electron transport in photosystem I | chloroplast | 0,6 | 0,7 | 0,5 | 0,9 |
| 13 | Phvul.002G304400.1 | AT5G56840 | MYB-like transcription factor family protein | | transcription | nucleus | 0,6 | 0,5 | 0,5 | 1,0 |
| 14 | Phvul.003G117000.1 | AT4G16270 | Peroxidase superfamily protein | | unknown | extracellular region | 1,2 | 1,3 | 0,5 | 1,3 |
| 15 | Phvul.011G009700.1 | AT1G51400 | Photosystem II 5 kD protein | | response to wounding | chloroplast | 0,6 | 0,7 | 0,5 | 0,9 |
| 16 | Phvul.001G191600.1 | AT2G36190 | Cell wall invertase 4 | | sucrose catabolic process | extracellular region | 1,7 | 0,8 | 0,5 | 2,8 |
| 17 | Phvul.009G033500.1 | AT1G72230 | Cupredoxin superfamily protein | | xylan biosynthetic process | plasma membrane | 0,3 | 1,1 | 0,4 | 1,1 |
| 18 | Phvul.001G248700.1 | AT5G58380 | SOS3-interacting protein 1 | | signal transduction | plasma membrane | 1,0 | 1,0 | 0,3 | 1,1 |
| 19 | Phvul.007G156000.1 | AT4G28780 | GDSL-like Lipase/Acylhydrolase superfamily protein | | stomatal complex morphogenesis | extracellular region | 2,4 | 4,6 | 0,3 | 1,0 |
| 20 | Phvul.008G041200.1 | AT5G15230 | GAST1 protein homolog 4 | | gibberellic acid mediated signaling | extracellular region | 0,6 | 1,0 | 0,3 | 1,0 |
| 21 | Phvul.009G128500.1 | AT1G77380 | Aino acid permease 3 | | negative regulation of apoptosis | plasma membrane | 1,5 | 2,0 | 0,2 | 0,9 |
| 22 | Phvul.010G153700.1 | AT3G15680 | Ran BP2/NZF zinc finger-like superfamily protein | | plant-type cell wall organization | unknown | 1,2 | 1,1 | DOWN | 3,3 |
| 23 | Phvul.011G058900.1 | Solyc10g051330 | Receptor-like kinase | | signal transduction | extracellular region | 5,3 | 1,7 | DOWN | 1,2 |
| 24 | Phvul.009G156300.1 | AT5G64530 | Xylem NAC domain 1 | | regulation of apoptosis | nucleus | 4,9 | 3,2 | DOWN | 1,2 |
| 25 | Phvul.011G044900.1 | AT4G39250 | RAD-like 1 | | transcription | nucleus | DOWN | 5,4 | DOWN | 1,4 |
| 26 | Phvul.011G209700.1 | Solyc03g095317 | Quinone oxidoreductase-like protein 2 | | Zinc-binding dehydrogenase | extracellular region | 3,5 | 6,6 | DOWN | 1,0 |
| 27 | Phvul.011G206100.1 | AT5G07030 | Eukaryotic aspartyl protease family protein | 2 | proteolysis | cell wall | 9,5 | 10,3 | DOWN | 1,0 |
| 28 | Phvul.008G252000.1 | Solyc12g056800 | Oxidoreductase family protein | | unknown | extracellular region | 0,7 | 13,7 | DOWN | 2,0 |
| 29 | Phvul.008G087100.1 | AT2G38540 | Lipid transfer protein 1 | | lipid transport | chloroplast | 5,4 | 17,5 | DOWN | 1,2 |
| 30 | Phvul.001G025800.1 | AT5G59845 | Gibberellin-regulated family protein | | response to gibberellin | extracellular region | 9,8 | 18,6 | DOWN | 1,0 |
| 31 | Phvul.003G101600.1 | Solyc07g065490 | Protein DEK | | negative regulation of transcription | extracellular region | 41,2 | 21,6 | DOWN | 1,0 |
| 32 | Phvul.002G231400.1 | AT1G07400 | HSP20-like chaperones superfamily protein | | response to oxidative stress | cytoplasm | 138,3 | 112,1 | DOWN | 1,8 |

Colored cells are significantly different compared to the control treatment, adjusted p-value<0.05

**Table S9.** Bean genes whose expression is **up-regulated by Te but not by Tu**. Annotations are based on the refence loci from Arabidopsis or tomato. The 4 most-right columns show the fold change (FC) values of plants treated with salicylic acid (SA), jasmonic acid (JA), *T. urticae* (Tu) or *T. evansi* (Te), relative to control. These genes were **selected based on** the criteria that expression is significantly up-regulated in the Te sample, but not in the Tu sample. This table is ranked according to the name in the 'involved in' column.

| | | | | | | FC expression, relative to control | | | |
|---|---|---|---|---|---|---|---|---|---|
| **#** | **Bean locus ID** | **REFERENCE** | **ANNOTATION** | **INVOLVED IN** | **LOCALIZATION** | **SA** | **JA** | **Tu** | **Te** |
| 1 | Phvul.004G136000.1 | AT1G73190 | Aquaporin-like superfamily protein | autophagy | central vacuole | NA | UP | NA | UP |
| 2 | Phvul.004G108600.1 | AT4G25720 | Glutaminyl cyclase | negative regulation of defense | chloroplast | NA | UP | NA | UP |
| 3 | Phvul.011G204600.1 | Solyc06g035940 | Homeobox leucine zipper protein | unknown | nucleus | NA | UP | NA | UP |
| 4 | Phvul.011G129800.1 | Solyc03g116860 | Acyl-CoA N-acyltransferases superfamily protein | unknown | cytoplasm | UP | UP | NA | UP |
| 5 | Phvul.001G098600.1 | AT1G52140 | Avr9/Cf-9 rapidly elicited protein | unknown | extracellular region | UP | UP | NA | UP |
| 6 | Phvul.010G106500.1 | AT2G46240 | BCL-2-associated athanogene 6 | response to stress | nucleus | UP | UP | NA | UP |
| 7 | Phvul.007G018700.1 | AT5G05320 | FAD/NAD(P)-binding oxidoreductase family protein | response to stress | unknown | UP | UP | NA | UP |
| 8 | Phvul.007G199000.1 | AT5G27000 | Kinesin 4 | protein binding | cytoplasm | UP | UP | NA | UP |
| 9 | Phvul.010G127500.1 | AT3G42880 | Leucine-rich repeat protein kinase family protein | protein phosphorylation | extracellular region | UP | UP | NA | UP |
| 10 | Phvul.006G069500.1 | AT2G30020 | Protein phosphatase 2C family protein | defense response | plastid | UP | UP | NA | UP |
| 11 | Phvul.007G275900.1 | Solyc05g012850 | Transmembrane protein | unknown | membrane | UP | UP | NA | UP |
| 12 | Phvul.003G288200.1 | AT2G22590 | UDP-Glycosyltransferase superfamily protein | unknown | unknown | UP | UP | NA | UP |

Colored cells are significantly different compared to the control treatment, adjusted p-value<0.05

**Table S10.** Bean genes whose expression is **down-regulated by Te but not by Tu**. Annotations are based on the refence loci from Arabidopsis or tomato. The 4 most-right columns show the fold change (FC) values of plants treated with salicylic acid (SA), jasmonic acid (JA), *T. urticae* (Tu) or *T. evansi* (Te), relative to control. These genes were **selected based on** the criteria that expression is significantly down-regulated in by Te, but not by Tu. This table is ranked according to the name in the 'involved in' column.

| | | | | | | FC expression, relative to control | | | |
|---|---|---|---|---|---|---|---|---|---|
| **#** | **Bean locus ID** | **REFERENCE** | **ANNOTATION** | **INVOLVED IN** | **LOCALIZATION** | **SA** | **JA** | **Tu** | **Te** |
| 1 | Phvul.001G173300.1 | AT5G02790 | Glutathione S-transferase family protein | glucosinolate biosynthetic process | cytoplasm | 89,6 | 121,1 | 1,7 | DOWN |
| 2 | Phvul.011G204200.1 | Solyc09g031650 | RING/U-box superfamily protein | response to zinc ion | extracellular region | 1,2 | 45,5 | 1,0 | DOWN |
| 3 | Phvul.001G084000.1 | AT5G61890 | Integrase-type DNA-binding superfamily protein | defense response | nucleus | 33,6 | 25,6 | 1,2 | DOWN |
| 4 | Phvul.008G070000.1 | AT2G01770 | Vacuolar iron transporter 1 | intracellular sequestering of iron ion | vacuolar membrane | 5,9 | 17,9 | 1,1 | DOWN |
| 5 | Phvul.008G053400.1 | AT1G05840 | Eukaryotic aspartyl protease family protein | cell death | unknown | 24,9 | 12,1 | 1,0 | DOWN |
| 6 | Phvul.005G171800.1 | AT3G15990 | Sulfate transporter 3;4 | unknown | plasmodesma | 24,3 | 9,1 | 2,2 | DOWN |
| 7 | Phvul.002G135700.1 | Solyc05g053320 | Retrovirus-related Pol polyprotein from transposon TNT 1-94 | RNA-directed DNA polymerase activity | extracellular region | 2,0 | 4,7 | 1,5 | DOWN |
| 8 | Phvul.011G130200.1 | Solyc06g073940 | Protein FAF | negative regulation of phosphorylation | chloroplast | 2,8 | 3,9 | 1,4 | DOWN |
| 9 | Phvul.011G142600.1 | AT5G23960 | Terpene synthase 21 | sesquiterpene biosynthetic process | chloroplast | DOWN | 3,3 | 1,3 | DOWN |
| 10 | Phvul.010G073500.1 | AT5G01300 | Phosphatidylethanolamine-binding protein | myo-inositol hexakisphosphate biosynthesis | cytoplasm | 2,4 | 2,7 | 1,5 | DOWN |
| 11 | Phvul.008G108000.1 | AT2G38060 | Phosphate transporter 4;2 | nitrate transport | plastid | 2,3 | 2,3 | 3,4 | DOWN |
| 12 | Phvul.007G043000.1 | AT3G19550 | Glutamate racemase | unknown | mitochondrion | 4,3 | 2,1 | 1,2 | DOWN |
| 13 | Phvul.002G042400.1 | AT3G22600 | Bifunctional inhibitor/lipid-transfer protein | lipid transport | plasma membrane | 1,8 | 1,8 | 1,3 | DOWN |
| 14 | Phvul.009G008900.1 | AT3G18830 | Polyol/monosaccharide transporter 5 | negative regulation of defense | plasma membrane | 2,2 | 1,7 | 1,5 | DOWN |
| 15 | Phvul.011G065500.1 | AT1G07645 | Dessication-induced 1VOC superfamily protein | unknown | extracellular region | 4,0 | 1,7 | 1,0 | DOWN |
| 16 | Phvul.006G085100.1 | AT3G02710 | ARM repeat superfamily protein | D-xylose metabolic process | nucleus | 1,6 | 1,3 | 1,1 | 0,9 |
| 17 | Phvul.007G000900.1 | AT5G13150 | Exocyst subunit exo70 family protein C1 | pollen tube growth | exocyst | 2,8 | 1,0 | 0,9 | DOWN |
| 18 | Phvul.002G055600.1 | AT5G60900 | Receptor-like protein kinase 1 | defense response | plasma membrane | 1,0 | 0,9 | 6,9 | DOWN |
| 19 | Phvul.011G173600.1 | AT2G40720 | Tetratricopeptide repeat (TPR)-like superfamily protein | unknown | mitochondrion | 1,3 | 0,8 | 1,0 | 0,5 |
| 20 | Phvul.010G005900.1 | AT3G09220 | Laccase 7 | response to nitrate | extracellular region | 1,2 | 0,8 | 2,9 | DOWN |
| 21 | Phvul.001G169700.1 | AT2G37530 | Forkhead box protein G1 | unknown | mitochondrion | 0,8 | 0,8 | 2,7 | DOWN |
| 22 | Phvul.004G019000.1 | AT3G01420 | Peroxidase superfamily protein | cell death | extracellular region | 1,9 | DOWN | 8,8 | DOWN |
| 23 | Phvul.001G087500.1 | AT5G14020 | Endosomal targeting BRO1-like domain-containing protein | unknown | vacuole | 1,8 | DOWN | 1,0 | DOWN |

Colored cells are significantly different compared to the control treatment, adjusted p-value<0.05

**Table S11.** Bean genes whose expression is **down-regulated by both Te and Tu** (19/19 down) Annotations are based on the refence loci from Arabidopsis or tomato. The 4 most-right columns show the fold change (FC) values of plants treated with salicylic acid (SA), jasmonic acid (JA), *T. urticae* (Tu) or *T. evansi* (Te), relative to control. These genes were **selected based on** the criterium that expression is significantly down-regulated in both the Te sample and the Tu sample. This table is ranked according to the name in the 'involved in' column.

| | | | | | | FC expression, relative to control | | | |
|---|---|---|---|---|---|---|---|---|---|
| # | Bean locus ID | REFERENCE | ANNOTATION | INVOLVED IN | LOCALIZATION | SA | JA | Tu | Te |
| 1 | Phvul.011G166900.1 | AT5G24090 | Acidic chitinase | defense response | extracellular region | DOWN | 50,3 | DOWN | DOWN |
| 2 | Phvul.004G129700.1 | AT1G17860 | Kunitz family trypsin and protease inhibitor | response to stress | extracellular region | 1,0 | 31,4 | DOWN | DOWN |
| 3 | Phvul.010G165000.1 | AT1G16760 | Protein kinase | plant-type cell wall modification | nucleus | DOWN | 17,6 | DOWN | DOWN |
| 4 | Phvul.011G209800.1 | Solyc02g086800 | Phospholipid-transporting ATPase | phospholipid translocation | plasma membrane | 3,2 | 15,7 | DOWN | DOWN |
| 5 | Phvul.001G238800.1 | AT3G55700 | UDP-Glycosyltransferase superfamily protein | transferring glycosyl groups | nucleus | 2,0 | 11,9 | DOWN | DOWN |
| 6 | Phvul.003G116900.1 | AT3G17130 | Plant invertase/pectin methylesterase inhibitor | unknown | extracellular region | 4,7 | 9,7 | DOWN | DOWN |
| 7 | Phvul.010G155300.1 | AT1G52560 | HSP20-like chaperones superfamily protein | response to stress | chloroplast | 3,2 | 6,6 | DOWN | DOWN |
| 8 | Phvul.010G146300.1 | AT4G32890 | GATA transcription factor 9 | circadian rhythm | nucleus | 1,4 | 6,3 | DOWN | DOWN |
| 9 | Phvul.001G212200.1 | AT4G02090 | Multidrug resistance protein ABC transporter | transmembrane transport | nucleus | 1,3 | 5,9 | DOWN | DOWN |
| 10 | Phvul.006G165900.1 | AT2G42800 | Receptor like protein 29 | signal transduction | plasma membrane | 1,5 | 5,8 | DOWN | DOWN |
| 11 | Phvul.003G230300.1 | AT1G32560 | Late embryogenesis abundant protein, group 1 | response to stress | plasmodesma | 7,4 | 4,9 | DOWN | DOWN |
| 12 | Phvul.007G086400.1 | AT5G48100 | Laccase/Diphenol oxidase family protein | flavonoid biosynthetisis | unknown | 0,7 | 1,6 | DOWN | DOWN |
| 13 | Phvul.003G212700.1 | AT5G52020 | Integrase-type DNA-binding protein | defense response | nucleus | 0,9 | 1,5 | DOWN | DOWN |
| 14 | Phvul.003G031300.1 | AT1G68530 | 3-ketoacyl-CoA synthase 6 | jasmonic acid biosynthesis | unknown | 2,9 | 1,5 | DOWN | DOWN |
| 15 | Phvul.003G027400.1 | AT1G13250 | Galacturonosyltransferase-like 3 | carbohydrate biosynthetisis | Golgi apparatus | 0,8 | 1,5 | 0,3 | 0,5 |
| 16 | Phvul.005G001100.1 | AT1G71691 | GDSL-like Lipase/Acylhydrolase protein | unknown | extracellular region | 2,1 | 1,2 | DOWN | DOWN |
| 17 | Phvul.002G233600.1 | AT5G37020 | Auxin response factor 8 | organ morphogenesis | nucleus | 1,0 | 1,0 | 0,5 | 0,7 |
| 18 | Phvul.003G196300.1 | AT3G48100 | Response regulator 5 | signal transduction | nucleus | 0,8 | 0,5 | 0,3 | 0,6 |
| 19 | Phvul.009G096900.1 | AT1G52820 | 2-oxoglutarate (2OG) and Fe(II)-dependent oxygenase | unknown | cytoplasm | 0,2 | 0,4 | DOWN | DOWN |

Colored cells are significantly different compared to the control treatment, adjusted p-value<0.05

**Table S12.** Bean genes whose expression is **up-regulated by both Te.** Annotations are based on the refence loci from Arabidopsis or tomato. The 5 most-right columns show the fold change (FC) values of plants treated with salicylic acid (SA), jasmonic acid (JA), *T. urticae* (Tu) or *T. evansi* (Te), relative to control or *T. urticae* relative to *T. evansi* (Tu/Te). These genes were **selected based on** the criterium that expression is significantly up-regulated in both the Te sample and the Tu sample. This table is ranked according to the expression of *T. urticae*, relative to *T. evansi*.

#### Table S12

| | | | | | | | FC expression, relative to control | | | | |
|---|---|---|---|---|---|---|---|---|---|---|---|
| # | Bean locus ID | REFERENCE | ANNOTATION | LOCI | INVOLVED IN | LOCALIZATION | SA | JA | Tu | Te | Tu/Te |
| 1 | Phvul.003G128200.1 | AT1G19250 | Flavin-dependent monooxygenase 1 | | defense response | chloroplast | UP | NA | UP* | UP | 730,9 |
| 2 | Phvul.007G049700.1 | AT4G21410 | Cysteine-rich RLK (RECEPTOR-like protein kinase) 29 | | response to abscisic acid | plasma membrane | UP | UP | UP* | UP | 510,1 |
| 3 | Phvul.002G038700.1 | AT5G13930 | Chalcone and stilbene synthase family protein | 3 | flavonoid biosynthesis | vacuole | UP | UP | UP* | UP | 442,0 |
| 4 | Phvul.010G120200.1 | AT5G54160 | O-methyltransferase 1 | 2 | phenylpropanoid metabolism | cytoplasm | UP | UP | UP* | UP | 313,9 |
| 5 | Phvul.008G287200.1 | AT1G59960 | NAD(P)-linked oxidoreductase superfamily protein | | response to stress | cytoplasm | UP | UP | UP* | UP | 280,3 |
| 6 | Phvul.009G244000.1 | AT4G37340 | Cytochrome P450, family 81, subfamily D, polypeptide 3 | | oxygen binding | unknown | UP | UP | UP* | UP | 256,0 |
| 7 | Phvul.006G129500.1 | AT5G06730 | Peroxidase superfamily protein | | response to oxidative stress | vacuolar membrane | UP | NA | UP | UP | 172,8 |

*Table continues on next page*

**Table S12** *continued*

| # | Bean locus ID | REFERENCE | ANNOTATION | LOCI | INVOLVED IN | LOCALIZATION | FC expression, relative to control | | | | |
|---|---|---|---|---|---|---|---|---|---|---|---|
| | | | | | | | SA | JA | Tu | Te | Tu/Te |
| 8 | Phvul.002G134600.1 | AT4G16070 | Mono-/di-acylglycerol lipase class 3 | | lipid metabolic process | plasma membrane | UP | NA | UP* | UP | 95,1 |
| 9 | Phvul.003G192000.1 | AT5G15130 | WRKY DNA-binding protein 72 | | defense response | nucleus | UP | UP | UP* | UP | 78,0 |
| 10 | Phvul.006G124800.1 | AT1G30700 | FAD-binding Berberine family protein | | toxin catabolic process | chloroplast | UP | UP | UP* | UP | 77,6 |
| 11 | Phvul.005G054100.1 | AT1G78380 | Glutathione S-transferase TAU 19 | 2 | toxin catabolic process | chloroplast | UP | UP | UP* | UP | 69,8 |
| 12 | Phvul.005G054100.1 | AT1G78380 | Glutathione S-transferase TAU 19 | 2 | toxin catabolic process | chloroplast | UP | UP | UP* | UP | 69,8 |
| 13 | Phvul.007G011300.1 | AT2G37430 | C2H2 and C2HC zinc fingers superfamily protein | | response to stress | nucleus | UP | UP | UP* | UP | 68,6 |
| 14 | Phvul.007G227900.1 | AT2G18950 | Homogentisate phytyltransferase 1 | 2 | defense response | plasma membrane | UP | NA | UP* | UP | 60,9 |
| 15 | Phvul.005G080300.1 | AT3G56400 | WRKY DNA-binding protein 70 | 2 | defense response | nucleus | UP | UP | UP* | UP | 57,8 |
| 16 | Phvul.006G039800.1 | AT4G31500 | Cytochrome P450, family 83, subfamily B, polypeptide 1 | | response to stress | membrane | NA | UP | UP* | UP | 53,1 |
| 17 | Phvul.001G040600.1 | AT5G10530 | L-type lectin receptor kinase IX.1 | 2 | defense response | extracellular region | UP | NA | UP* | UP | 48,1 |
| 18 | Phvul.002G076500.1 | AT1G70170 | Matrix metalloproteinase | | proteolysis | extracellular region | UP | UP | UP* | UP | 40,4 |
| 19 | Phvul.006G194000.1 | AT4G27280 | Calcium-binding EF-hand family protein | | response to wounding | plasma membrane | UP | UP | UP | UP | 33,1 |
| 20 | Phvul.005G162700.1 | Solyc11g022470 | MYB family transcription factor family protein | | signal transduction | nucleus | NA | NA | UP* | UP | 22,0 |
| 21 | Phvul.006G056600.1 | Solyc01g086810 | NBS-LRR disease resistance protein | | defense response | extracellular region | UP | UP | UP* | UP | 18,7 |
| 22 | Phvul.008G098500.1 | AT5G05600 | 2-oxoglutarate (2OG) and Fe(II)-dependent oxygenase | 2 | response to jasmonic acid | cytoplasm | UP | UP | UP | UP | 18,4 |
| 23 | Phvul.004G099300.1 | AT2G34930 | Disease resistance family protein / LRR family protein | 2 | defense response | cell wall | UP | NA | UP | UP | 17,7 |
| 24 | Phvul.001G019200.1 | AT4G37260 | MYB domain protein 73 | | response to jasmonic acid | nucleus | UP | UP | UP* | UP | 17,2 |
| 25 | Phvul.004G125400.1 | AT5G62200 | Embryo-specific protein 3, (ATS3) | | response to wounding | vacuolar membrane | UP | NA | UP* | UP | 17,2 |
| 26 | Phvul.006G033300.1 | AT5G60900 | Receptor-like protein kinase 1 | 2 | defense response | plasma membrane | UP | UP | UP | UP | 15,0 |
| 27 | Phvul.008G249900.1 | AT5G05340 | Peroxidase superfamily protein | | protein binding | cel wall | UP | NA | UP* | UP | 13,8 |
| 28 | Phvul.001G042100.1 | AT1G80840 | WRKY DNA-binding protein 40 | | defense response | nucleus | 90,3 | 14,2 | 816* | 69,2 | 11,8 |
| 29 | Phvul.010G013600.1 | AT1G47890 | Receptor like protein 7 | | defense response | unknown | UP | NA | UP* | UP | 9,8 |
| 30 | Phvul.007G024800.1 | AT2G39210 | Major facilitator superfamily protein | | defense response | chloroplast | UP | UP | UP* | UP | 9,7 |
| 31 | Phvul.011G189300.1 | AT1G76680 | 12-oxophytodienoate reductase 1 | | response to wounding | cytoplasm | UP | UP | UP* | UP | 9,4 |
| 32 | Phvul.011G056100.1 | AT5G44640 | Beta glucosidase 13 | | glucosinolate catabolism | Golgi apparatus | UP | UP | UP | UP | 8,8 |
| 33 | Phvul.001G076700.1 | AT2G44480 | Beta glucosidase 17 | | response to stress | extracellular region | UP | UP | UP | UP | 7,3 |
| 34 | Phvul.007G125500.1 | AT2G38870 | Serine protease inhibitor, potato inhibitor type I | | defense response | extracellular region | UP | UP | UP | UP | 6,4 |
| 35 | Phvul.010G055100.1 | AT5G48770 | Disease resistance protein (TIR-NBS-LRR class) family | | defense response | cytoplasm | UP | UP | UP | UP | 4,6 |
| 36 | Phvul.001G095600.1 | AT1G76650 | Calmodulin-like 38 | | response to wounding | nucleus | UP | UP | UP* | UP | 4,5 |
| 37 | Phvul.011G109600.1 | AT3G23250 | MYB domain protein 15 | | signal transduction | nucleus | UP | UP | UP | UP | 3,9 |
| 38 | Phvul.011G056700.1 | AT1G20480 | AMP-dependent synthetase and ligase family protein | | phenylpropanoid metabolism | unknown | UP | NA | UP | UP | 3,9 |
| 39 | Phvul.008G011600.1 | AT3G12580 | Heat shock protein 70 | | response to stress | cytoplasm | UP | UP | UP | UP | 3,8 |
| 40 | Phvul.008G058400.1 | AT3G61510 | ACC synthase 1 | | ethylene biosynthesis | cytoplasm | UP | UP | UP | UP | 3,6 |
| 41 | Phvul.011G182300.1 | AT4G08850 | Leucine-rich repeat receptor-like protein kinase | | response to stress | membrane | UP | UP | UP | UP | 3,6 |
| 42 | Phvul.011G189200.1 | AT1G76690 | 12-oxophytodienoate reductase 2 | | response to wounding | cytoplasm | UP | UP | UP | UP | 2,7 |
| 43 | Phvul.008G250600.1 | AT4G35160 | O-methyltransferase family protein | | phenylpropanoid metabolism | cytoplasm | UP | UP | UP | UP | 2,6 |
| 44 | Phvul.007G030800.1 | AT1G25390 | Protein kinase superfamily protein | 2 | defense response | plasma membrane | UP | UP | UP | UP | 2,1 |
| 45 | Phvul.005G026100.1 | AT5G42380 | Calmodulin like 37 | | ethylene biosynthesis | cytoplasm | UP | UP | UP | UP | 1,8 |
| 46 | Phvul.007G077800.1 | AT5G24090 | Acidic chitinase | | defense response | extracellular region | UP | UP | UP | UP | 1,6 |
| 47 | Phvul.009G046500.1 | AT4G27670 | Heat shock protein 21 | | response to stress | chloroplast | UP | UP | UP | UP | 0,8 |

Colored cells are significantly different compared to the control treatment, adjusted p-value<0.05
